# Molecular Structure and DNA Binding Mode of Unsymmetric Cyanine Dyes RiboGreen and OliGreen

**DOI:** 10.64898/2026.05.04.722657

**Authors:** Nolan Blackford, Saileena Nepal, Huan He, Lianqing Zheng, Wei Yang, Robert Silvers

**Affiliations:** Department of Chemistry & Biochemistry, Florida State University, Tallahassee, FL 32306, USA; Institute of Molecular Biophysics, Florida State University, Tallahassee, FL 32306, USA; College of Medicine, Translational Science Laboratory, Florida State University, Tallahassee, FL 32306, USA

**Keywords:** Nucleic acid dyes, nucleic acid binding, RiboGreen, OliGreen, PicoGreen

## Abstract

The binding of fluorescent dyes to nucleic acids and their fluorogenic properties are indispensable tools for nucleic acid detection, quantification, and imaging, yet the molecular structures of several widely used commercial dyes have remained unknown. Here, we *de novo* determined the molecular structures of RiboGreen and OliGreen and confirmed the previously proposed structure of PicoGreen using high-field NMR spectroscopy and ultra-high resolution mass spectrometry. All three dyes were identified as unsymmetric cyanine dyes, where benzazolium and 4-quinolinium moieties are linked by a monomethine bridge. Complete ^1^H and ^13^C resonance assignments enabled us to expand the existing chemical shift reference set for this important class of dyes. Photophysical characterization with standardized single- and double-stranded DNA and RNA targets indicated that all dyes performed similarly upon binding despite being marketed towards different nucleic acid types. NMR spectroscopy and long-timescale molecular dynamics simulations showed that RiboGreen interacts with double-stranded DNA predominantly by two binding modes, electrostatic interactions with the phosphodiester backbone and π-π stacking with accessible nucleobases of the DNA molecule. These results establish the molecular structures of three widely used commercial dyes and provide a structural and mechanistic framework for understanding the fluorogenic properties of this class of dyes.

**Highlights:**

- Determination of the molecular structures of nucleic acid dyes RiboGreen, OliGreen, and PicoGreen
- NMR spectroscopic characterization of all three dyes.
- NMR and MD data indicate binding to be dominated by electrostatic and π-π stacking interactions

## INTRODUCTION

Fluorescent dyes that bind to DNA and RNA are indispensable for the study of nucleic acids *in vitro* and *in vivo* alike. Among the different classes of fluorophores, unsymmetric cyanine dyes are the most widely used, commercially available dyes employed in a wide range of applications involving nucleic acid detection. Common uses include nucleic acid quantification[1–3], visualization and staining in gel electrophoresis,[4–11] visualization *in vivo* and *ex vivo* using fluorescence microscopy,[12–16] and nucleic acid amplification and hybridization assays.[17–24] These dyes have also enabled more specialized, “off-label” applications. For example, thiazole orange (TO) has been used in fluorescent intercalator displacement assays to probe ligand binding in G-quadruplex DNA,[25, 26] and its capacity to bind folded i-motif DNA has been exploited to monitor i-motif folding.[27] Likewise, OliGreen (OG) and PicoGreen (PG) have been used in aptamer biosensor applications.[28–32] More recently, RiboGreen (RG) enabled direct observation of RNA stability and RNA ligand binding using a highly sensitive and readily parallelizable approach called differential scanning fluorimetry on RNA,[33, 34] and has enabled the precise quantification of nucleic acid cargo in nanoparticles used in RNA drug formulations.[35, 36]

Unfortunately, only limited information – particularly regarding their molecular structure – is provided by manufacturers for several of these key commercial dyes, which can be problematic for applications where understanding of binding properties matter. While many variations for unsymmetric cyanine dyes exist, we here focus on some of the most commonly used, commercially available, green-fluorescent, and monomeric dyes (Figure 1). Previous studies have determined the molecular structures of SYBR Gold (SGO),[37] SYBR Green I and II (SGI/SGII),[38] proposed the structure of PG,[23] and have shown that all are derivatives of TO or oxazole yellow (YO), consisting of benzazolium and 4-quinolinium moieties connected via a monomethine bridge. Figure 1 depicts the general scaffold and chemical constitution of common commercial unsymmetric cyanine dyes. These dyes are differentiated by the identity of four R groups. The substitution found on N^1^ (R^1^) is typically hydrophobic (either aliphatic or aromatic) in nature, e.g. a methyl, propyl, butyl, or phenyl group. Most R^2^ groups – often dubbed the “tail” – are hydrocarbon chains ending in strongly basic tertiary amines which are protonated under physiological conditions or quaternary amines bearing a permanent positive charge. R^6^ and R^7^ are typically hydrogen, with the notable exceptions of SGO and OG that carry a methoxy moiety at R^6^ and R^7^, respectively.

**Figure 1:**
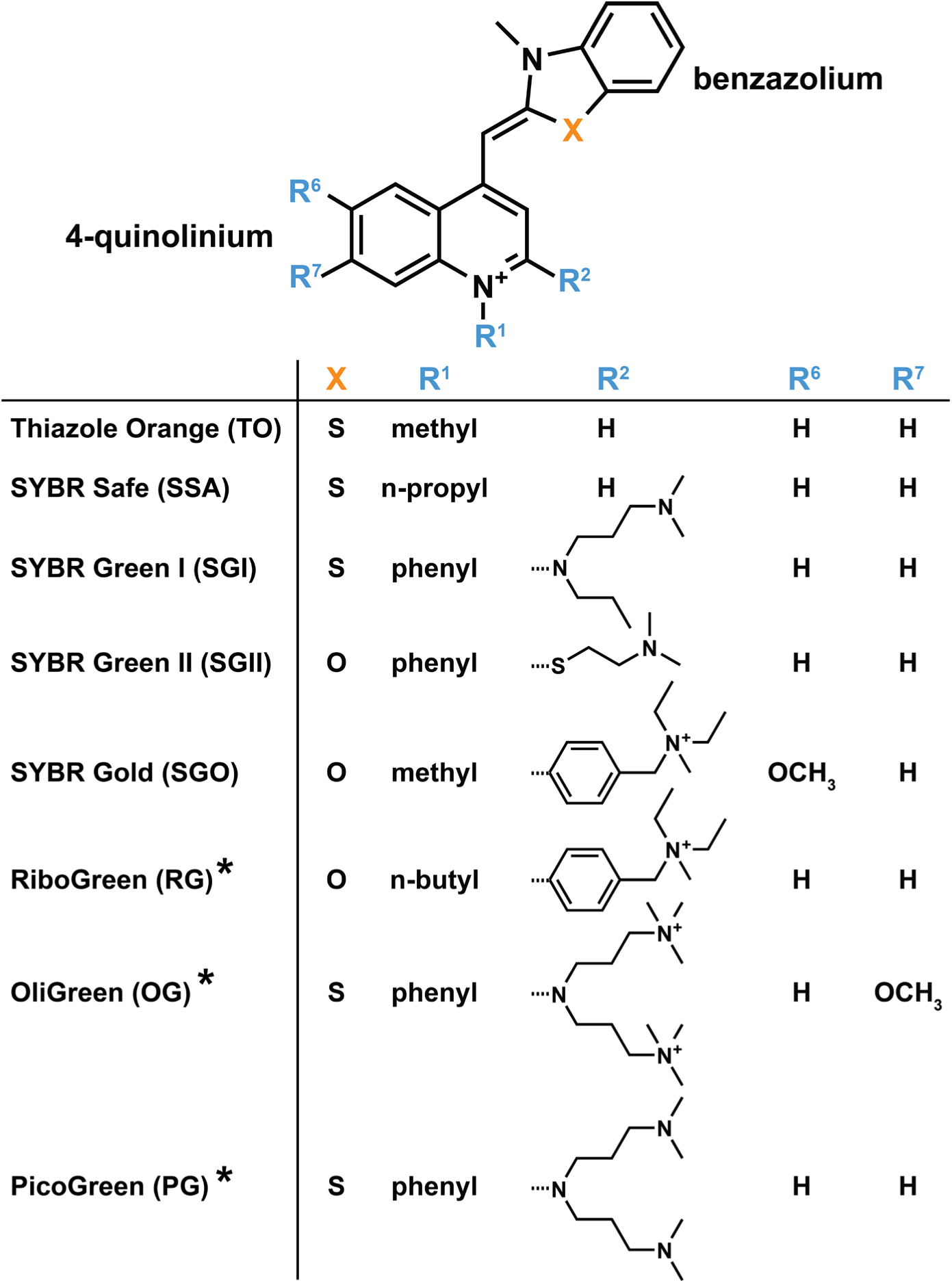
Chemical constitution of common commercial unsymmetric cyanine dyes derived from thiazole orange (TO) and oxazole yellow (YO). The dyes all share a common scaffold (top) and are differentiated by the identity of R groups attached to the 4-quinolinium moiety as well as the identity of X in the benzazolium moiety (X=O or X=S). Positively charged “tails” and/or aromatic R groups contribute to the dye’s affinity toward nucleic acids by creating additional sites for electrostatic interactions and π-π stacking interactions, respectively. The numbering of the R groups is based on the numbering proposed previously for this class of dyes.[37, 39] The asterisk (*) indicates dyes with molecular structures determined in this work.

RG and OG are widely used and commercially available nucleic acid binding dyes with previously unknown molecular structure. While it has been suggested that they belong to the class of unsymmetric cyanine dyes, this was not known for certain. RG is marketed specifically for use in RNA quantification, and OG is marketed for use in single-stranded DNA quantification, but both have been used “off-label” for applications including differential scanning fluorimetry on RNA.[33, 34] It is unclear if the difference in marketing is indicative of a difference in structure, binding mode, or fluorescent properties of these dyes which differentiate them from the others with known structures.

Here, we *de novo* determined the molecular structures of RG and OG using high-field NMR spectroscopy and ultra-high resolution mass spectrometry (MS), as well as investigated the DNA binding mode of RG using NMR spectroscopy and long-timescale molecular dynamics (MD) simulations. Furthermore, we confirmed the molecular structure of PG whose structure was proposed previously based on comparable photophysical and binding properties as well as mass spectrometry.[38] Data presented here close an important gap in the mechanistic understanding of these three dyes. NMR data and long-timescale MD simulations provide a mechanistic understanding of dominant interactions between dye and double-stranded DNA. Overall, this work expands the molecular foundation for more informed use of RG, OG, and PG in biochemical, biophysical, and analytical applications. It also contributes to the larger goal of connecting fluorophore structure to binding mechanisms and fluorogenic properties, an essential step toward the rational selection and future design of nucleic acid dyes for specific experimental settings.

## MATERIALS AND METHODS

### NMR Spectroscopy

The fluorescent dyes RiboGreen (RG), OliGreen (OG), and PicoGreen (PG) were ordered from Thermo Fisher Scientific as 1 mL stocks suspended in DMSO. Each dye was prepared for NMR spectroscopy by first removing the non-deuterated DMSO by lyophilizing the entire 1 mL stock solution in the absence of light using a Labconco Free Zone Benchtop Freeze Dryer. After ∼36 hours, the dried dye samples were resuspended in ∼100 µL of DMSO-d_6_ (Cambridge Isotope Laboratories) and lyophilized for 24 additional hours to reduce the amount of lingering non-deuterated DMSO in the final NMR sample. The dried dyes were then each resuspended in 400 µL of DMSO-d_6_ and transferred into 5 mm Shigemi NMR tubes. Because NMR spectra of PG were broadened in DMSO-d_6_, the PG samples were freeze dried again and the solvent was exchanged to methanol-d_4_, which led to a dramatic improvement in line width. NMR experiments were performed on a 700 MHz Bruker Avance III spectrometer equipped with a TCI triple resonance cryo-probe. For each dye, a suite of experiments consisting of [^1^H]-1D, two-dimensional [^1^H, ^1^H]-DQF-COSY, [^1^H, ^1^H]-TOCSY,[40] [^1^H, ^1^H]-NOESY,[41, 42], [^1^H, ^13^C]-HSQC,[43–45] and long-range [^1^H, ^13^C]-HMBC spectra were acquired. For NMR spectroscopy on the 10-mer A+B duplex DNA, experiments with jump-return echo sequences were used for observing imino protons at 279 K. A [^1^H]-1D and [^1^H, ^1^H]-NOESY (mixing time of 250 ms) were collected on the DNA before and after the addition of freeze-dried RG dye.

### Mass Spectrometry

RG and OG were analyzed by direct infusion mass spectrometry (MS), while PG was analyzed by liquid chromatography mass spectrometry (LC-MS) due to the presence of impurities. RG and OG samples were diluted to ∼ 30 μM in 90% aqueous methanol and were ionized by nano-electrospray ionization (nESI) and detected by an Orbitrap Exploris 480 mass spectrometer (Thermo Fisher Scientific, Waltham, MA).

The ionization voltage was at 2.2 kV and ion transfer tube temperature of 275°C. The precursor ions were detected with a mass resolution of 120 k, Automatic Gain Control (AGC) of 3 million ions and a maximum injection time of 50 ms. Most abundant precursor ions were molecular ions and were selected for tandem MS with a 40% higher-energy collisional dissociation (HCD) collision energy, mass resolution of 120 k, isolation window of 2 Da, and a maximum injection time of 100 ms. The PG sample was diluted to ∼3 μM in 10% aqueous methanol with 0.1% formic acid (FA). 18 uL of the solution was injected via sample loop onto a liquid chromatography Dionex UltiMate 3000 system (Thermo Fisher Scientific, Waltham, MA) equipped with an Accucore C18 analytical column (100 x 2.1 mm, Thermo Fisher Scientific, Waltham, MA). The mobile phases were A (99.9% H_2_O, 0.1% FA) and B (99.9% acetonitrile, 0.1% FA). The gradient profile is as follows: hold at 0% B for 1 min; then a linear gradient to 98% B over 20 min; then hold at 98% B for 10 min; then a linear gradient back to 1% B over 1 min; re-equilibrate the column at 1% B for 8 min with a flow rate of 200 μL/min. The eluate was ionized by electrospray ionization (ESI) with a heated ESI (HESI) source with a sheath gas flow rate of 3 and an auxiliary gas flow rate of 3 and a spray voltage of 3.9 kV. Ions were detected by the Q Exactive HF MS (Thermo Fisher Scientific, Waltham, MA) with high mass accuracy and high resolving poser. The precursor ions were detected with a m/z range of 100 – 1500 Da, a mass resolution of 120 k and an AGC target of 3 million ions. Then targeted tandem MS of PG molecular ions were carried out with an isolation window of 2 Da, mass resolution of 30k, AGC target of 1 million ions. Data was acquired and visualized by Xcalibur (Thermo Fisher Scientific, Waltham, MA).

### Preparation of DNA and RNA samples

Two sets of complimentary 10-mers (one RNA and one DNA) were ordered from Genscript Biotech, named 10-mer A and B with the following sequences:

DNA-A: 5’-AGCTCGCATG-3’
DNA-B: 5’-CATGCGAGCT-3’
RNA-A: 5’-AGCUCGCAUG-3’
RNA-B: 5’-CAUGCGAGCU-3’

Each was diluted to 100 µM with DEPC-treated water. For experiments on single-stranded DNA and RNA, 10-mer A was used. For experiments on double-stranded DNA and RNA, 10-mers A and B were duplexed by mixing equal quantities of each followed by heating to 90°C for 10 minutes and subsequent snap cooling on ice. To prepare the 10-mer A+B DNA for NMR spectroscopy, lyophilized 10-mer A and B were suspended in 10 mM sodium phosphate buffer containing 10% D_2_O and 0.01% DSS to a concentration of 200 µM and mixed in equal quantities for a final duplex concentration of 100 µM. For the addition of RiboGreen dye to the duplex NMR sample, 150 µL of RiboGreen stock solution was lyophilized and added to the NMR sample.

### UV-Vis and Fluorescence Spectroscopy

The extinction coefficient of unbound OG was determined by first determining the molar concentration of our working stock using quantitative NMR with maleic acid as standard as shown previously.[46] Briefly, DMSO was removed by freeze-drying a specified volume of OG dissolved in DMSO and subsequent resuspension in the same volume of TE buffer (10 mM Tris pH 7.5 and 1 mM EDTA). Subsequently, OG concentrations of 5.4, 2.7, 1.35, 0.675, and 0.3375 μM were prepared by dilution in TE buffer. Absorbance spectra were recorded using an Agilent Cary 300 series UV-Vis spectrophotometer with a scan rate of 600 nm/min, a data interval of 1 nm and an averaging time of 0.1 seconds. The intensity at the absorbance maximum at a wavelength of 494 nm was plotted against the dye concentration. A linear fit was used to determine the molar extinction coefficient per Beer’s law. Furthermore, samples of each dye, both free and bound to ssDNA, dsDNA, ssRNA, and dsRNA were prepared. Each sample contained 5 µg/mL of nucleic acid (excluding dye only samples) and 200-fold diluted dye in 10 mM Tris pH 7.5, 1 mM EDTA and 2% DMSO. Dye concentrations were measured in dilution factor from the stock concentration as the manufacturer does not disclose concentration. The same samples were used for both UV-Vis and fluorescence spectroscopy. UV-Vis spectroscopy was used to measure the maximum absorption wavelength of each dye both free and when bound to each nucleic acid type using an Agilent Cary 300 series UV-Vis spectrophotometer. Measurements were performed in a 10 mm sub micro cell quartz cuvette (part no: 6610024100). Absorbance scans were performed at room temperature from 800 to 200 nm with an averaging time of 0.1 seconds at 600 nm/min and a data interval of 1 nm. A fluorescence spectrophotometer (Cary Eclipse fluorescence spectrophotometer Model G98001) was used to measure the excitation and emission wavelengths of each sample. The absorption maxima obtained from the UV-VIS spectrophotometer was used as the excitation wavelength for the emission scan. We then used the emission maximum from the emission scan to perform the excitation scan. Runs were carried out at the slowest scan rate of 30 nm/min with a PMT detector voltage of 600V.

### Dissociation Constant Determination using McGhee-von Hippel Model

Double-stranded 10-mer DNA and RNA were prepared by mixing 10-mers A and B of each type in equal quantities. They were then refolded by heating to 90°C for 10 minutes then snap cooled on ice. From this, 20 nucleic acid concentrations were prepared via serial dilutions in TE buffer. Each condition was evaluated at three dye concentrations (1.0, 0.5, and 0.1 µM) for RG, OG, and PG. The concentrations of the dyes were determined by using their molar extinction coefficients (PG: ε_500nm_ ≈ 70,000 M^-1^cm^-1^,[1] RG: ε_482nm_ ≈ 67,000 M^-1^cm^-1^,[2] and OG: ε_494nm_ ≈ 77,000 M^-1^cm^-1^ (determined here)). The samples of all sets were loaded onto a black 384-well plate (Nunc® MaxiSorp™) and fluorescence was measured using a TECAN Spark® microplate reader. Measurements were recorded with an excitation wavelength of 500 nm, an emission wavelength of 525 nm, a bandwidth of 10 nm, and 100 flashes per well. The gain of the instrument was set to “high dynamic range” mode. The McGhee-von Hippel model was employed to fit the data as shown previously for SGO.[37, 47] Briefly, the following fitting equation was used

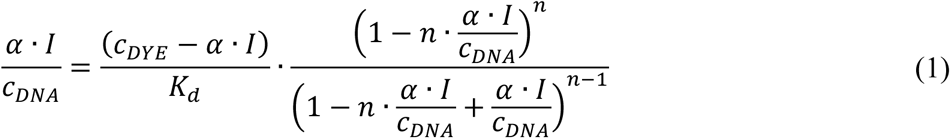

where *I* are the measured fluorescence intensities at varying total DNA concentrations *c_DNA_*, *K_d_* is the dissociation constant, n is the binding site size, and *c_DYE_* is the total dye concentration. α is a proportionality constant. *I*, *c_DNA_*, and *c_DYE_* are experimentally known. For each set of data consisting of titration curves at three different dye concentrations, a global fit was performed where *K_d_* and n were global fitting parameters and α was a local fitting parameter.

### Molecular Dynamics Simulations

The binding of RG to dsDNA was investigated using molecular dynamics (MD) simulations. The initial setups of all the simulations were prepared using the CHARMM-GUI platform.[48] The initial length of the simulation box was set to 80 Å. Potassium and chloride ions were added at a concentration of 0.15 M to neutralize the system. The CHARMM force field was used to treat the DNA, dyes, and ions,[49, 50], while the TIP3P model was applied for the water molecules.[51] The Particle Mesh Ewald (PME) algorithm was used to calculate the long-range electrostatic interactions.[52] The binding of RG to the DNA was performed using the Orthogonal Space Tempering (OST) method[53–55] with the distance between the mass centers of the dye and the DNA as the order parameter. For the production simulations, an NPT ensemble was applied with a reference temperature of 300 K and pressure of 1 atm.[56] All the simulations were performed using a modified version of the CHARMM[57] program developed by the Yang group.

## RESULTS AND DISCUSSION

### Molecular Structures of RG, OG, and PG

The molecular structures of RG and OG are currently unknown, while the structure of PG was previously postulated based on comparable photophysical and binding properties as well as mass spectrometry.[38] Here, we *de novo* determined the structures of RG and OG, as well as confirmed the structure of PG by high-field NMR spectroscopy and ultra-high resolution mass spectrometry using established approaches (Figure 2).[37–39] A suite of experiments consisting of [^1^H]-1D, [^1^H, ^1^H]-DQF-COSY, [^1^H, ^1^H]-TOCSY, [^1^H, ^1^H]-NOESY, [^1^H, ^13^C]-HSQC, and long-range [^1^H, ^13^C]-HMBC spectra were employed to fully assign all ^1^H and ^13^C carbon chemical shifts and determine the molecular structures of RG (Figures S1-S5), OG (Figures S6-S10), and PG (Figures S11-S18). Furthermore, the molecular structures were analyzed using ultra-high resolution MS to determine total masses and fragmentation patterns (Figures S19-S24). The observed and expected *m/z* ratios of the monoisotopic species of [RG]^2+^ (m/z_obs_=253.661, m/z_calc_=253.662), [OG]^3+^ (m/z_obs_=204.124, m/z_calc_=204.124), and [PG]^+^ (m/z_obs_=552.315, m/z_calc_=552.317) are in excellent agreement. For PG, a second protonated species was detected [PG+H]^2+^ (m/z_obs_=276.661, m/z_calc_=276.661). The characteristic isotopic distribution confirms the presence of exactly one sulfur atom in OG and PG, but not RG (Figures S19B, S21B, and S23). In MS/MS spectra, tertiary and quaternary amines lead to characteristic patterns due to fragmentation and neutral loss of the dye-specific amine moiety.[58] In OG, we observed the characteristic fragmentation pattern corresponding to the loss of trimethylamine at m/z 58.06507 (C_3_H_8_N^+^), m/z 59.07289 (C_3_H_9_N^•+^), and m/z 60.08075 (C_3_H_10_N^+^) (Figure S22). Furthermore, the neutral loss of trimethylamine (59.0735 Da) lead to a characteristic peak of [OG-N(CH_3_)_3_]^3+^ at m/z 184.43257. For RG, we observed similar patterns that can be ascribed to the fragmentation of the *N,N*-diethylmethylamine moiety at m/z 86.09640 (C_5_H_12_N^+^) and m/z 87.10524 (C_5_H_13_N^•+^) (Figure S20). A secondary fragmentation product was observed at m/z 72.08075 that can be attributed to the decomposition of *N,N*-diethylmethylamine to diethylamine. Additionally, the remaining dye fragment can be observed at m/z 420.21825. For PG, we employed LC-MS due to the presence of impurities and the presence of multiple protonation states (Figure S23). For both [PG]^+^ and [PG+H]^2+^ precursors, we observed fragmentation corresponding to the loss of *N,N*-dimethylpropylamine at m/z 86.09695 (C_5_H_12_N^+^) (Figure S24). Furthermore, the core structure formed by the decomposition of the entire tail is observed at m/z 367.12602.

**Figure 2:**
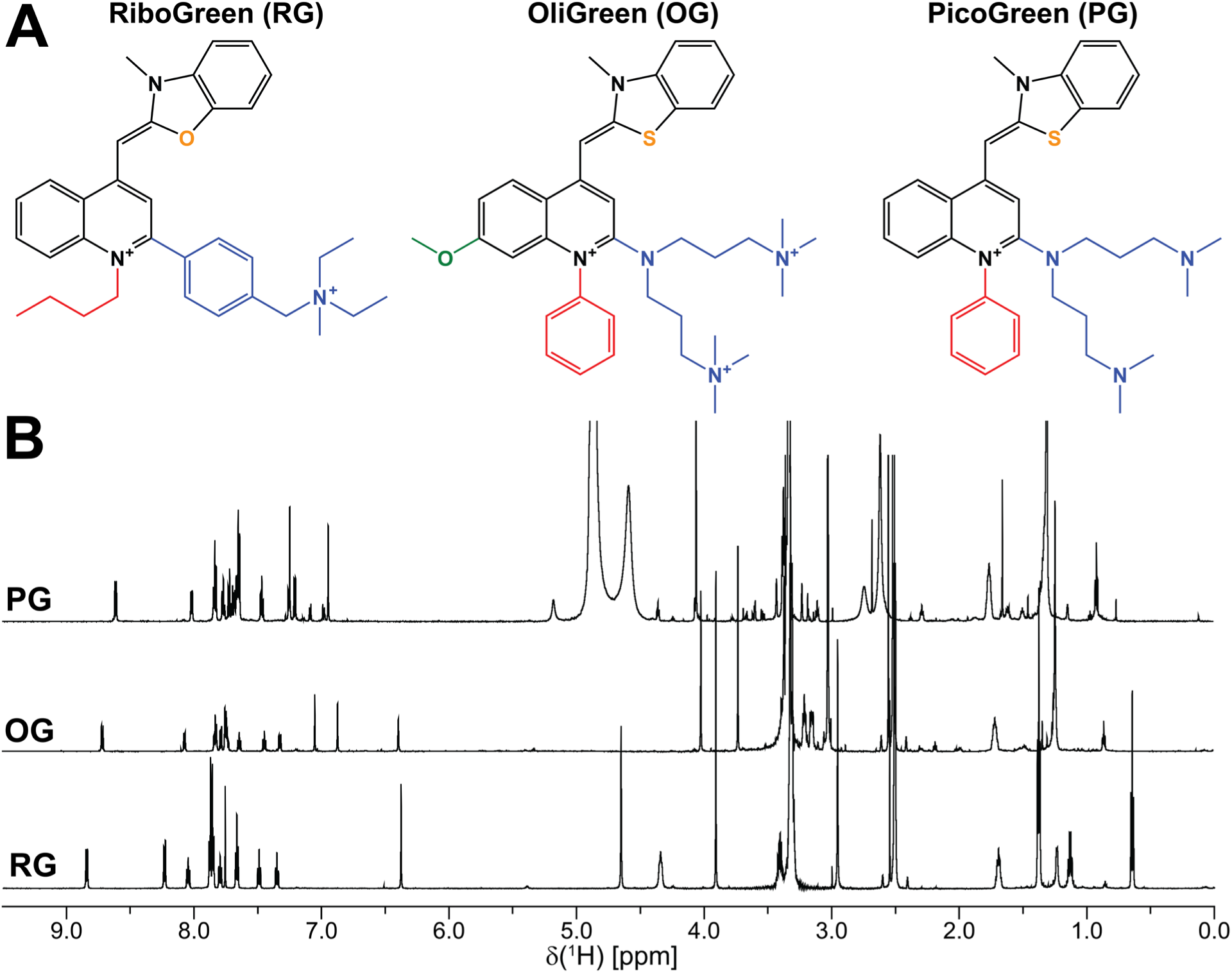
Molecular structures and 1D ^1^H NMR spectra of RG, OG, and PG. (A) Molecular structures of RG, OG, and PG. Oxygen or sulfur atoms within the benzazolium moiety are shown in orange. The substitution (R^1^) found on N^1^ is shown in red. The substitution (R^2^) – often referred to as the “tail” – found on C^2^ is shown in blue. The methoxy moiety present in OG on R^7^ is shown in green. (B) The spectra of RG and OG were recorded in DMSO-d_6_, the spectrum of PG was recorded in methanol-d_4_. Strong signals have their intensities truncated for the sake of clarity. All spectra were recorded at a field strength of 16.4 T.

The combination of NMR spectroscopy and MS confirmed that all three dyes share the same underlying scaffold as other unsymmetric cyanine dyes. RG’s scaffold is comprised of benz*<u>oxa</u>*zolium and 4-quinolinium moieties connected by a monomethine bridge, while OG’s and PG’s scaffolds are comprised of benzo*<u>thia</u>*zolium and 4-quinolinium moieties connected by a monomethine bridge. RG carries an n-butyl moiety at N^1^, while OG and PG both carry phenyl groups. On C^2^, RG is substituted by a rigid 4-((diethyl(methyl)ammonio)methyl)phenyl moiety identical to SGO, while PG and OG carry 2-(bis(3-(dimethylamino)propyl)amino and 2-(bis(3-(trimethylammonio)propyl)amino moieties, respectively. Besides carrying quaternary amines in its “tail”, OG also has one other distinctive feature compared to PG, a methoxy moiety in position R^7^.

### NMR Spectroscopy of Unsymmetric Cyanine dyes

The ^1^H and ^13^C chemical shifts for RG, OG, and PG were assigned in their entirety using high-field NMR spectroscopy (Tables 1 and S1). This new chemical shift data set in combination with chemical shift data reported previously for TO, SSA, SGI, SGII, and SGO,[37, 39] enabled the identification of characteristic chemical shift values indicative of certain structural elements and motifs which will be beneficial for future NMR spectroscopic studies on similar compounds. Substitutions on the benzazolium moiety are uncommon, thus chemical shift variations in this part of the molecule are small and can be used for referencing (Figure 3). In fact, the chemical shift footprint of this moiety is easily recognizable and the only chemical shift variations stem from the identity of the 1’ heteroatom which is either oxygen or sulfur. While the ^1^H and ^13^C chemical shifts of 2’, 3’-CH_3_, 4’, 5’, and 6’ are unperturbed by the presence of sulfur or oxygen, the chemical shifts of 3a’, 7a’, and 7’ are highly sensitive to the identity of X and can indeed be used to reliably determine if the dye carries a benz*<u>oxa</u>*zolium or a benzo*<u>thia</u>*zolium moiety (Figure 3A, green). Likewise, the ^1^H and ^13^C chemical shifts in the monomethine bridge (2a’) are highly sensitive to the identity of X. The chemical shift difference for X=S vs. X=O is most pronounced in 7a’ with over 20 ppm (^13^C), followed by 2a’ with 0.6/13.5 ppm (^1^H/^13^C), 7’ with 0.3/11.7 ppm (^1^H/^13^C), and 3a’ with 9.0 ppm (^13^C) (Figure 3C, orange vs. blue circles). Surprisingly, the chemical shift of the directly bonded 2’ carbon is virtually unperturbed, likely due to its location in between X and N^3’^ as well as its participation in photoisomerization. No long-range effects of the identity of X were observed in the 4-quinolinium moiety and chemical shifts are unaffected by X=S vs. X=O. However, with all R group substitutions situated in the 4-quinolinium moiety, chemical shifts in this part of the molecule are less predictable and are spread out as indicated by large errors (Figure 3C, open circles), except for ^13^C chemical shifts of quaternary carbons 4, 4a, and 8a. The ^1^H and ^13^C chemical shifts of 2 and 3 are affected by the presence and identity of R^2^, while the chemical shifts of 5, 6, 7, and 8 are affected by the presence and identities of R^6^ and R^7^ (Table S1).

**Table 1:**
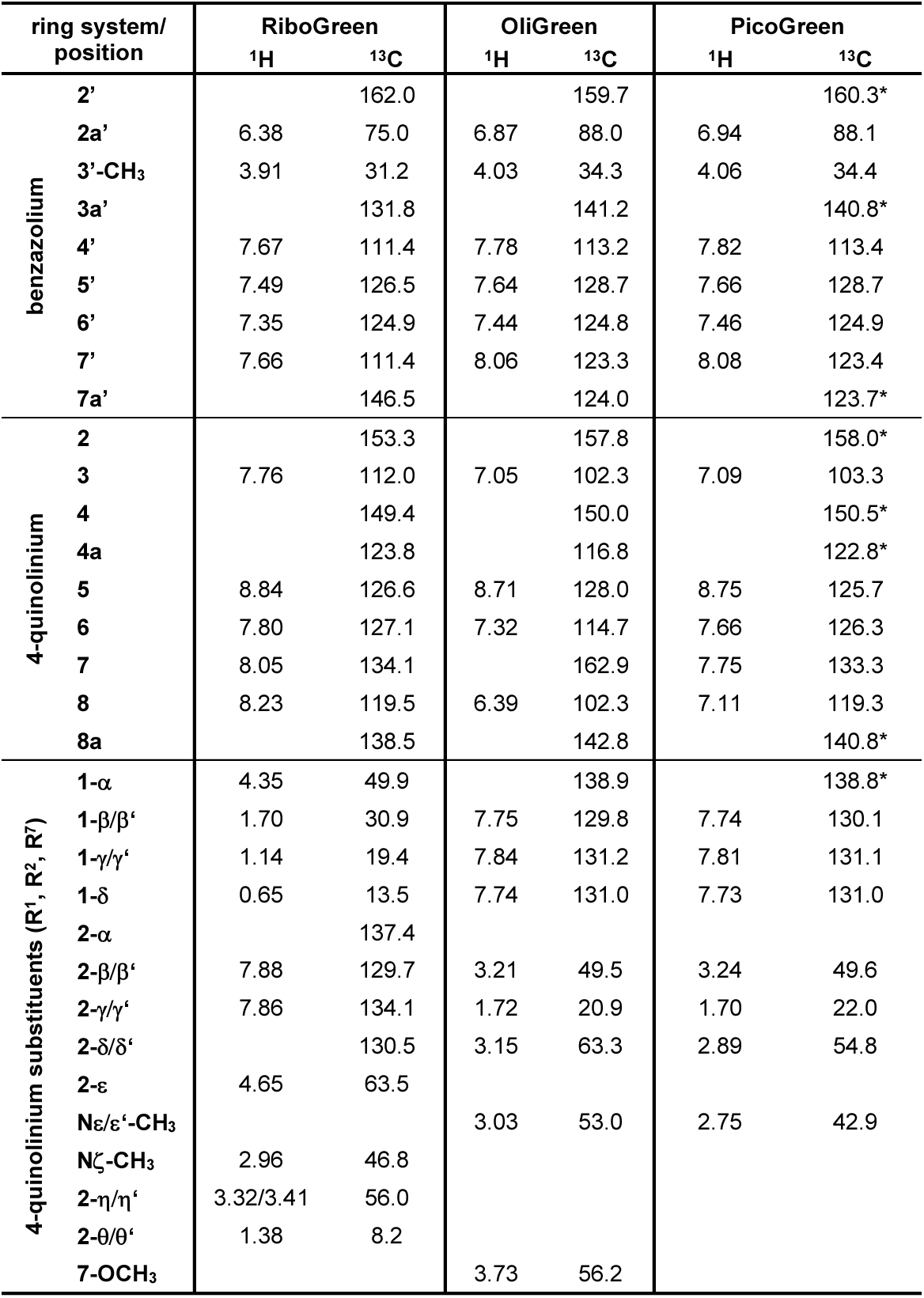
^1^H and ^13^C chemical shifts of RG, OG, and PG as determined by NMR spectroscopy. Atom naming in the scaffold and substituents follows convention previously proposed for this class of dyes.[37, 39]. All chemical shifts are from samples dissolved in DMSO-d_6_ except for certain PG ^13^C chemical shifts denoted with and asterisk (*), these signals were only visible in methanol-d_4_. An expanded table of ^1^H and ^13^C chemical shifts which include dyes from previous NMR studies in addition to the ones listed here can be found in Table S1. The numbering of all atoms can be found in Figure S25.

**Figure 3:**
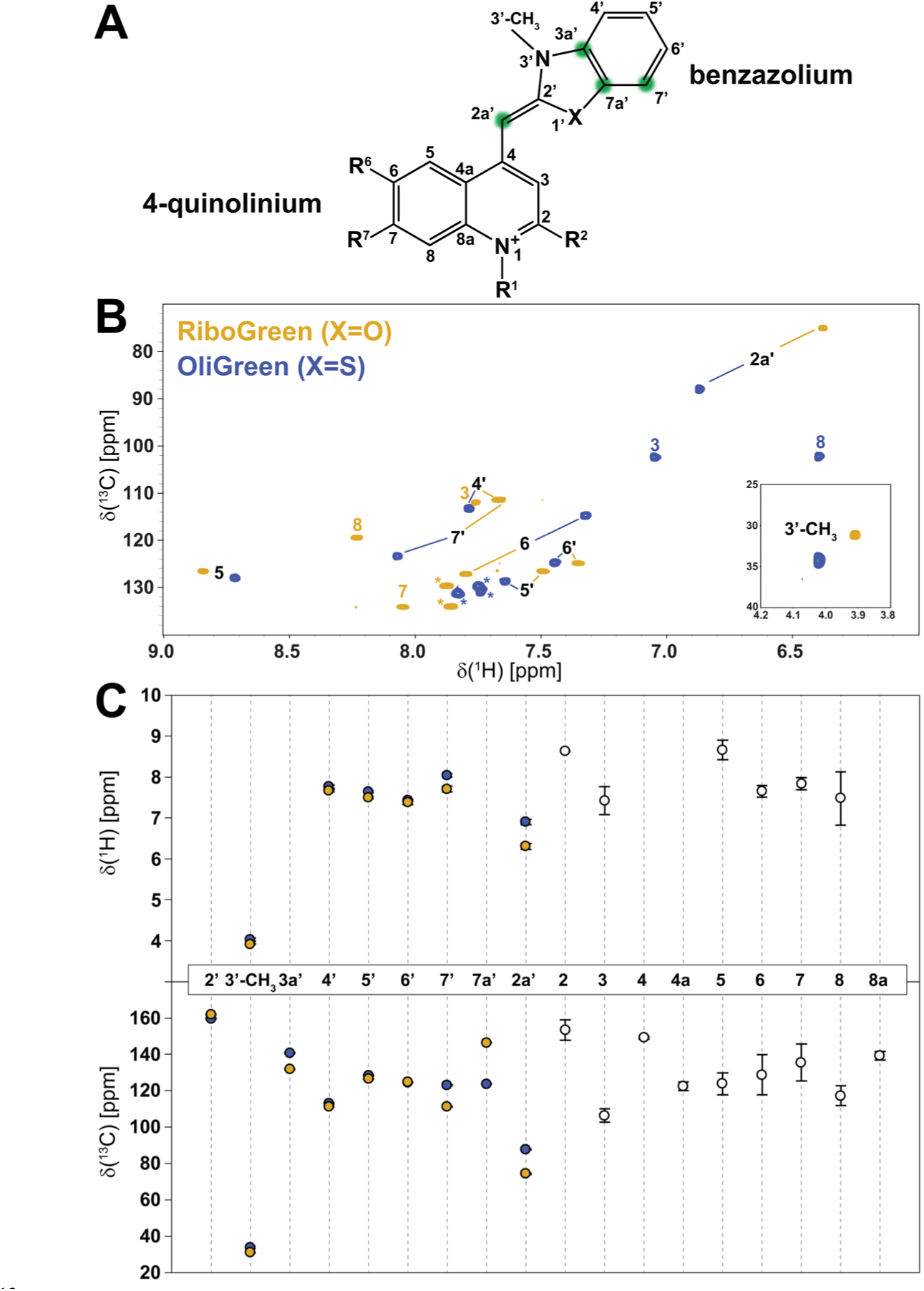
^1^H and ^13^C chemical shifts of dye scaffolds. (A) Dye scaffold including atom numbering. Locations in the benzazolium moiety highlighted in green are most susceptible to the identity of X (O vs. S). (B) An overlay of the ^1^H-^13^C HSQC aromatic regions of RG, which contains a benzoxazolium moiety, and OG, which contains a benzothiazolium moiety. Asterisks (*) indicate aromatic, non-scaffold moieties. (C) Plots of the average ^1^H and ^13^C chemical shifts at each position in the scaffold listed in Table S1 with error bars indicating the standard deviation. Within the benzazolium moiety, average chemical shifts of the three benzoxazoles (orange) and five benzothiazoles (blue) are shown, while average chemical shifts for the 4-quinolinium moiety (white) are based on all eight chemical shift data sets.

### Photophysical properties of RG, OG, and PG

Unsymmetric cyanine dyes possess large extinction coefficients and exhibit negligible fluorescence in the unbound state. The molar extinction coefficients of unbound RG and PG were determined previously as ε_482nm_ ≈ 67,000 M^-1^cm^-1^ and ε_500nm_ ≈ 70,000 M^-1^cm^-1^, respectively.[1, 2] Here, we report the molar extinction coefficient of unbound OG as ε_494nm_ ≈ 77,000 M^-1^cm^-1^ (Figure S26). Upon binding to nucleic acids, their quantum yield increases dramatically. This so-called fluorogenic property is thought to be rooted in the restriction of intramolecular torsion around the monomethine bridge. Computational studies for TO and its analogues have shown that rotation about the monomethine bridge beyond an interplanar angle of around 60° results in a dark state that decays non-radiatively.[59] This fluorescence quenching mechanism known as twisted intramolecular charge transfer (TICT) depends on twisted dye conformations that promote intramolecular charge separation and is exhibited by e.g. cyanine, aminocoumarin, 1,8-naphthalimide, boron dipyrromethene, and rhodamine dyes.[60] Thus, restriction of torsion about the monomethine bridge in unsymmetric cyanine dyes by nucleic acid binding leads to a substantial increase in excited state lifetimes.[37, 61, 62] A mild increase in excited state lifetimes attributed to the decrease in torsion around the monomethine bridge has also been observed previously when the dyes PG and SGI were placed in a high-viscosity environment.[23, 62] As expected, we also observed a mild fluorescence enhancement when each of the three dyes was placed in a solution of 100% glycerol (Figure S27).

The general photophysical properties of RG, OG, and PG as well as related dyes have been studied extensively.[1, 2, 23, 28, 38, 62–65] The increase in quantum yield upon binding depends on nucleic acid type and composition and is in first approximation attributable to two major factors. The first is the enhancement in quantum yield each dye experiences upon binding, i.e. the intrinsic quantum yield of the bound state. The second is the strength of binding itself, as a stronger binding dye will even in the case of identical intrinsic quantum yields lead to stronger fluorescence as there will be a higher fraction of dye bound at any given time. Because fluorescence intensity depends on a variety of factors including binding affinity, intrinsic quantum yield, nucleic acid type and composition, as well as binding mode, the description of fluorescence exhibited by RG, OG, and PG was limited to a qualitative and relative analysis by recording excitation and emission spectra of RG, OG, and PG bound to different nucleic acid model systems (ssDNA, dsDNA, ssRNA, and dsRNA). This relative comparison is of particular interest considering that RG, OG, and PG are marketed towards quantification of RNA, ssDNA, and dsDNA, respectively. Surprisingly, RG, OG, and PG display remarkably similar preferences for nucleic acid type and species (DNA vs. RNA and single-stranded vs. double-stranded). As seen in Figures 4 and S28, all dyes tested here provide the strongest fluorescence when bound to dsDNA with a somewhat reduced fluorescence when bound to dsRNA (RG: ∼95%, OG: ∼66%, PG: ∼71%). Compared to their double-stranded counterparts, fluorescence is greatly reduced when bound to single-stranded species (ssDNA/ssRNA). Slight variations in the excitation and emission maxima were also observed for each dye when bound to the different nucleic acid types which are reported in Table S2.

**Figure 4:**
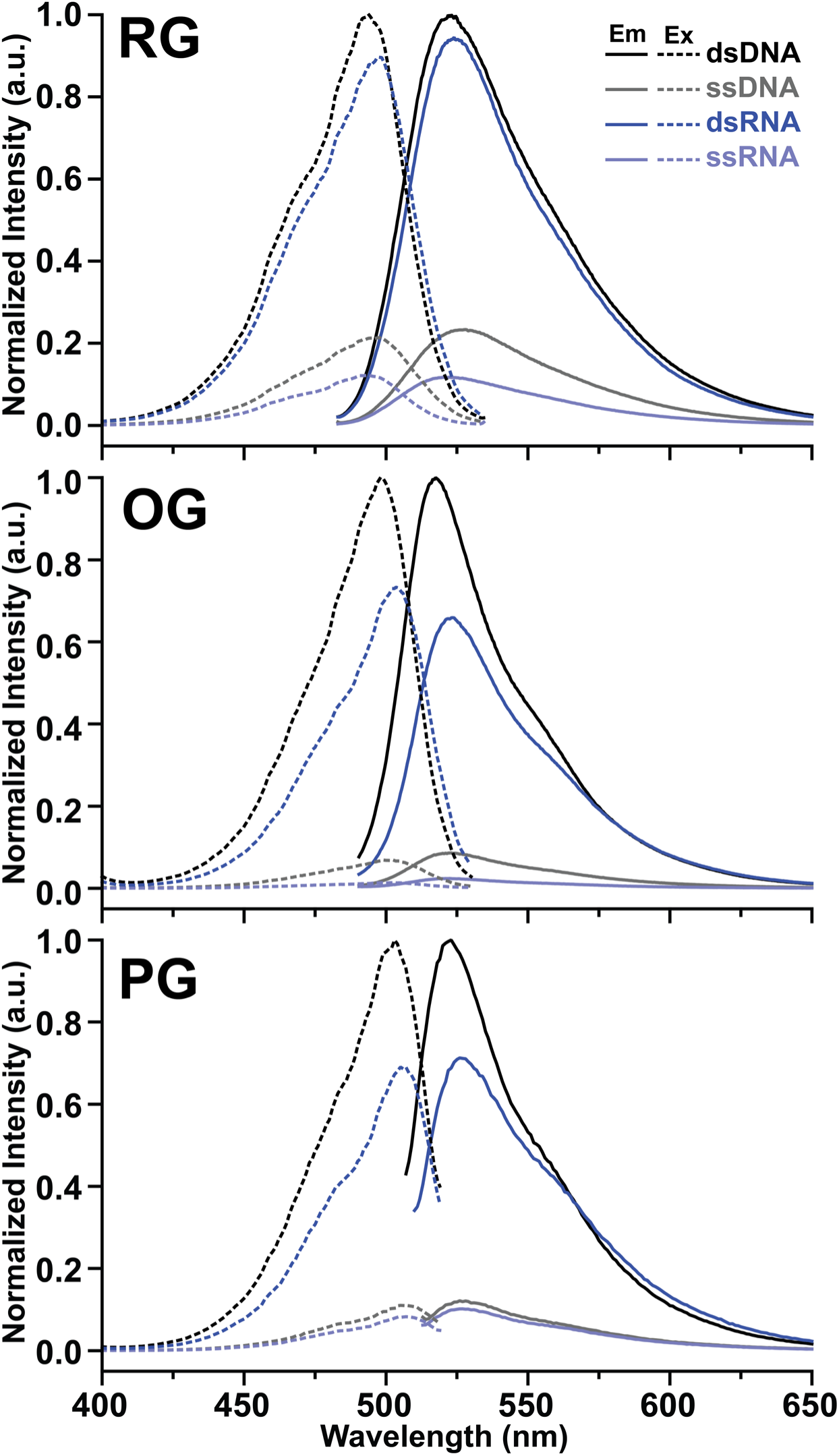
Emission and Excitation spectra of RG, OG, and PG. Emission and Excitation spectra of RG, OG, and PG bound to each type of nucleic acid, each comprised of the same 10-mer sequence (for single-stranded measurements 10-mer A was used). The plots are normalized to the largest intensities (Ex. or. Em.) produced by each dye. Dye concentrations and nucleic acid concentrations (measured in mg/mL) remain constant to enable relative comparisons between the dye’s fluorescent properties when bound to each nucleic acid type.

The similarities in relative fluorescence are significant considering the increasing number of applications where RG, OG, and PG are used “off label” (*vide supra*). It is also worth mentioning that the manufacturer notes that contamination with other nucleic acid species can affect the quantification due to cross-specificity, thus protocols recommend treating samples with appropriate DNases/RNases before running quantification assays when interfering contaminations are expected.

### Binding Affinities of RG, OG, and PG

While knowledge of the exact dye concentrations, binding affinities, and binding modes are not required when standard curves are employed in quantification assays, it is important to emphasize that the increasing use of these dyes for “off-label” applications require a more in-depth knowledge of their binding properties. We therefore determined the apparent dissociation constants of binding of RG, OG, and PG to 10-bp model dsDNA and dsRNA using the McGhee-von Hippel model (Table 2 and Figure S29).[37, 47, 66] Of the three dyes studied here, OG displayed the lowest apparent dissociation constant to both dsDNA and dsRNA of K_d_ ∼ 90-130 nM. This is likely due to the presence of three permanent positive charges in OG as well as a flexible tail that can accommodate stronger electrostatic interactions. In contrast, the binding to both dsDNA and dsRNA is slightly reduced in RG with a K_d_ ∼ 400-500 nM. This is expected considering that RG has only 2 positive charges and the “tail” is bulky and rigid in comparison to OG. PG displayed the largest apparent dissociation constant (K_d_ ∼ 0.8-1.0 μM), in line with previously reported dissociation constants.[66] The similarity in K_d_ between dsDNA and dsRNA for each dye indicated that the minor differences observed in fluorescence intensity (Figure 4) is likely due to small differences in quantum yield as opposed to a difference in binding strength. The binding sizes (n) observed in all dyes ranging from ∼ 3-4.5 are consistent with binding of a bulky dye molecule to a binding site that spans ∼ three base pairs in double-stranded nucleic acids.

**Table 2:** McGhee-von Hippel fitting parameters of the binding of RG, OG, and PG to dsDNA and dsRNA. Fitted graphs are shown in Figure S29.

| Dye | NA Type | Dissociation constant $K_d$ (nM) | Binding site size $n$ |
| --- | --- | --- | --- |
| RG | dsDNA | $371.0 \pm 102.7$ | $3.04 \pm 0.18$ |
| | dsRNA | $497.7 \pm 238.4$ | $3.00 \pm 0.33$ |
| OG | dsDNA | $129.2 \pm 81.7$ | $4.48 \pm 0.29$ |
| | dsRNA | $91.3 \pm 90.4$ | $4.52 \pm 0.41$ |
| PG | dsDNA | $1171.2 \pm 389.9$ | $2.34 \pm 0.33$ |
| | dsRNA | $757.9 \pm 431.8$ | $3.59 \pm 0.44$ |

### DNA Binding Mode of RG and OG

In the past, several binding modes have been identified by which ligands, including fluorescent dyes, interact with nucleic acids, namely interaction by (i) intercalation,[37, 38, 67] (ii) minor- or (iii) major-groove binding,[38, 68–72] (iv) external electrostatic or surface binding,[73–75] and (v) bis-intercalation.[76–78] Additionally, the interaction with the nucleic acid does not have to be limited to one single binding mode. Unsymmetric cyanine dyes have previously been hypothesized to have similar binding mechanisms, however, several studies into the binding mode of various dyes indicate a complex binding mode. For instance, single molecule magnetic tweezer experiments showed a lengthening and unwinding of the DNA helix upon binding to SGO in line with intercalation.[37] On the other hand, additional data obtained on the binding characteristics of several other dyes showed that the full picture is more complicated. For example, it was proposed that the positively charged tail present in many dyes extends into the minor groove and functions similarly to AT-hooks.[23, 38] Furthermore, computational studies have revealed binding mechanisms for several SGI/SGII related dyes which do not bind via intercalation, suggesting the possibility that not all unsymmetric cyanine dyes bind nucleic acid by the same mechanisms.[79] Binding is also affected by DNA:dye ratio, as circular dichroism studies on YO revealed a secondary dimeric binding mechanism, hypothesized to be surface binding as opposed to intercalation, which emerges when the ratio of dye to DNA is high.[80] For PG it was shown that while intercalation might dominate the interaction with double-stranded nucleic acid, PG was found to bind to single-stranded nucleic acids via a different mechanism with reduced affinity and quantum yield as well as showed a strong nucleotide preferences, preferring poly(dG) and poly(dT) over poly(dC) and poly(dA).[65]

To investigate the predominant binding mode of RG to double-stranded nucleic acids, we employed NMR spectroscopy and long-timescale MD simulations. Using a 10-bp dsDNA as model system, we investigated the effects of RG binding to DNA by monitoring changes in chemical shifts of imino protons involved in base pairing (Figure 5). In the absence of RG, a total of eight imino protons were observed, three stemming from H3 in thymidine and five from H1 in guanosine residues, all can be assigned using a NOESY spectrum (Figure 5, blue spectrum). Expectedly, the imino protons of the two terminal base pairs are broadened beyond detection due to the dynamics of the base pairs and the resulting rapid proton exchange with bulk water. Upon addition of sub-stoichiometric amounts of RG, a large number of new imino resonances appear in a concentration-dependent manner, while the original resonances reduce in intensity. Notably, many of these new resonances appear in the chemical shift ranges characteristic for terminal AT and GC base pairs around 11.5-12.5 ppm and 13.0-13.5 ppm, respectively (Figure 5, dashed boxes).[81–84] It is worth noting that these new resonances are also spread out, indicating that RG is not only interacting with the termini of duplex DNA, but also that this interaction cannot be uniformly ascribed to just one single state, but rather an array of different states. Furthermore, the appearance of additional resonances, some of which can be described as resonance splitting, indicated that the chemical environment of internal base pairs is affected as well by binding of RG. Overall, this is indicative of a complex binding mode of RG involving multiple internal and terminal binding sites that exist either simultaneously as a mixture and/or through exchange on the slow-exchange regime on the NMR timescale.

**Figure 5:**
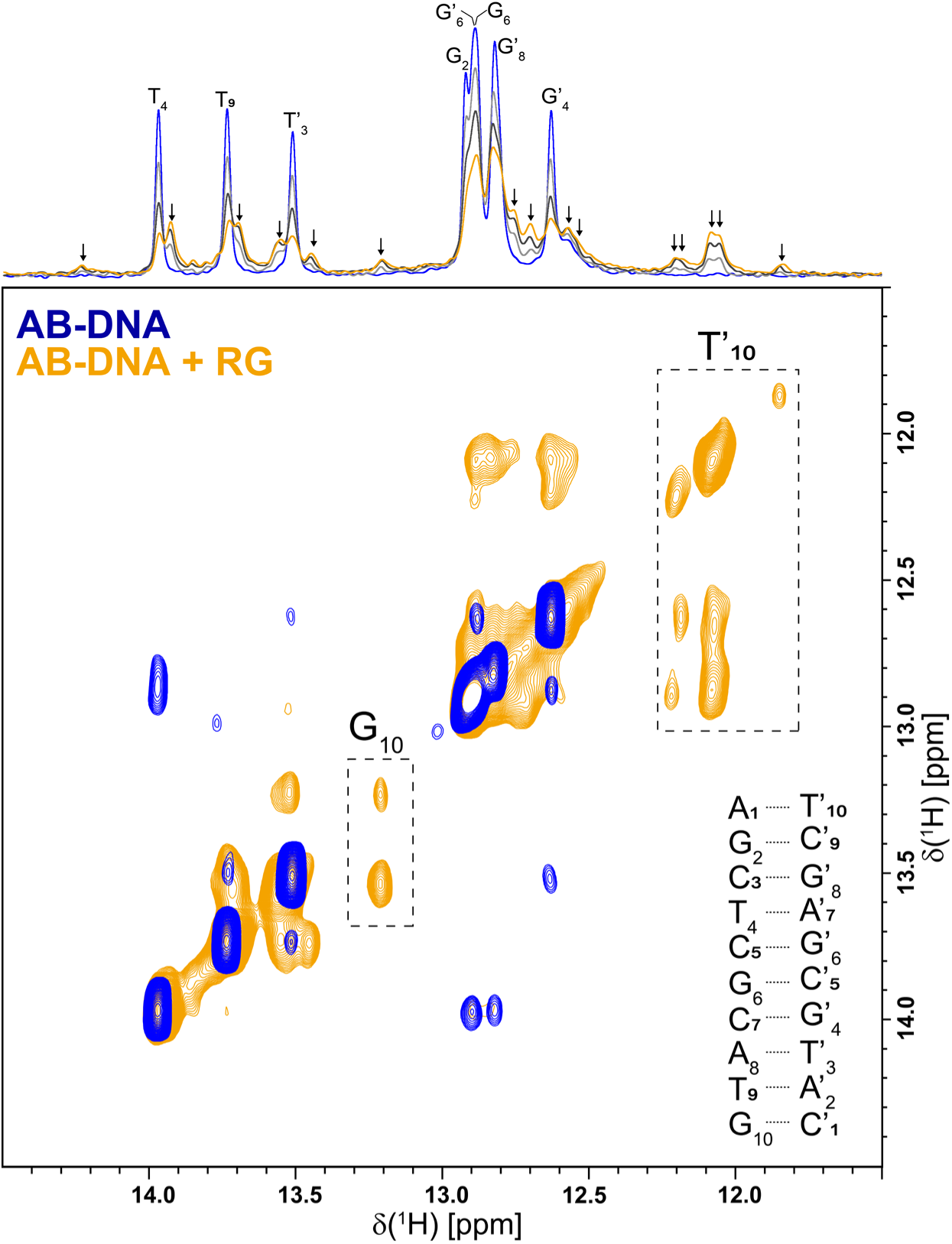
NMR spectroscopy of RG binding to dsDNA. Imino region of a [^1^H]-1D and 250 ms [^1^H, ^1^H]-NOESY that was acquired on a sample containing 100 µM A+B-DNA at 279 K and 700 MHz (blue). Resonance assignments are shown. 1D spectra of the titration of RG to the sample in three steps is shown (light gray, dark gray, orange), as well as the NOESY spectrum of the final titration step. New resonances in the 1D are indicated by arrows.

To resolve these binding sites, we employed long-timescale MD simulations that utilize the orthogonal space sampling scheme[53–55] to explore the slow-timescale motion-coupled binding of RG to dsDNA. In the orthogonal space tempering (OST) simulation, the distance between the centers of mass of the dsDNA (COM_DNA_) and RG (COM_Dye_) was employed as the order parameter. In the OST method, hidden degrees of freedom can be automatically accelerated, and thereby, random walks along the order parameter can be observed several times (Figure 6A, top). Such random walk behavior indicates the sufficiency of the sampling treatment. As a result, a converged free energy curve was obtained along the order parameter direction (Figure 6A, bottom). The results revealed that binding is driven predominantly by a combination of (i) electrostatic interactions between the positively charged RG dye and the negatively charged phosphodiester backbone of the DNA (Figure 6B, blue spheres) and (ii) π-π interactions between the aromatic rings systems of the dye and accessible nucleobases of the dsDNA (Figure 6B, orange spheres). The majority of these interactions with the DNA are very dynamic in nature and the dye samples the entire backbone and termini of the DNA indiscriminately of base identity. This is in line with observations from NMR spectroscopy that revealed a complex binding mode of RG involving multiple internal and terminal binding sites. Furthermore, this is also confirmed by the calculated free energy surface (Figure 6A, bottom), where the regions of binding free energy minima are shallow by less than 3 kcal/mol lower than the unbound region and COM distances (d_DNA-Dye_) range from ∼ 15 Å to ∼ 20 Å.

**Figure 6:**
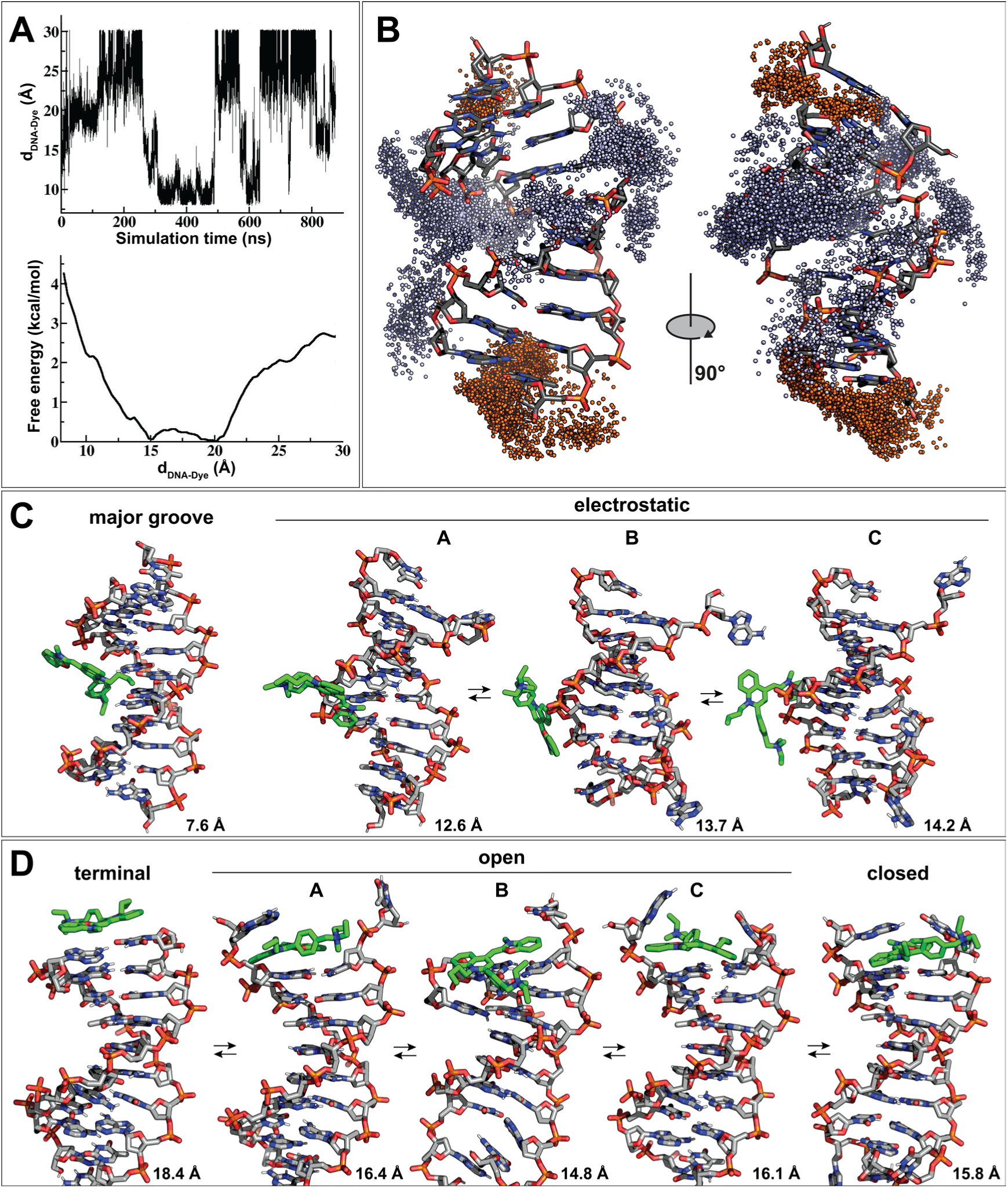
Molecular dynamics simulations of RG using OST. (A) (top) Fluctuation of the center of mass distance between the DNA and the dye with simulation time. The orthogonal space tempering method enables the sampling of both the binding and unbinding states of the dye to the DNA multiple times. (bottom) Free energy profile of the center of mass distance between the DNA and the dye. (B) Locations of bound dye molecules. Each sphere indicates the location of RG’s center of mass, when its center of mass is within 4 Å of any DNA heavy atom. Interactions with the backbone are dominated by electrostatics (blue spheres) and interactions with nucleobases of the DNA are dominated by π-π interactions (orange spheres). Interactions of RG (green) with (C) the major groove and backbone, as well as with (D) terminal and penultimate base pairs of the dsDNA. For each state, the distance between the centers of mass of dye and DNA is shown.

MD simulations revealed a variety of different binding modes as the dye dynamically samples the majority of the surface of the DNA molecule. Full, stable minor groove binding was not observed due to the bulkiness of the dye molecule and stable major groove binding (Figure 6C, *“major groove”*) was observed only rarely and the state was short-lived. The COM_DNA_-COM_Dye_ distance is small with 7.6 Å and the free energy is equal or higher to the free energy of the unbound state limit (∼3 kcal/mol). This is not surprising since RG has only very limited capabilities to form H-bonds with base pairs in the major groove. In most peripheral electrostatic interactions that were observed, the two positive charges in RG – one by the delocalized cyanine moiety and one by the quaternary nitrogen – align favorably with the backbone (Figure 6C). Even within the same general binding location, the dye rapidly samples various binding orientations (*“electrostatic” A, B, & C*). However, due to the bulkiness of the dye molecule, interactions with the minor groove are rare and short lived, while a substantial portion of the dye molecule tends to pivot towards the major groove instead. Notably, the oxygen atom of the benzoxazole ring forms a hydrogen bond with the phosphate in many of these interactions, while the benzoxazole ring itself forms hydrophobic contacts with the interface formed by H5/H6 of cytidine, H6 of thymidine, H8 of adenosine and guanosine residues as well as the H5/H5’ of the sugar moiety of the DNA.

RG also extensively interacts with the DNA through π-π interactions between the aromatic rings systems of the dye and accessible nucleobases of the dsDNA. In the *“terminal”* state, the dye molecule caps the end of dsDNA where the benzoxazolium/4-quinolinium moieties interact with the terminal base pairs (Figure 6D). It is common for the terminal base pairs to open and close, which is also observed during the MD simulation. When the *“open”* state is formed, the dye can access the penultimate base pairs and form π-π interactions with them. During this motion, the terminal bases of the DNA and dye sample many different states (*“open” A, B, &C*) with COM_DNA_-COM_Dye_ distances ranging from ∼ 15.0-16.5 Å. Notably, while internal intercalation events were not observed for RG, a partial and short-lived “intercalation” event can be observed when the previously flipped terminal bases partially close to form a weak base pair (*“closed”*). A similar behavior has been observed for MD simulations of ethidium bromide.[85]

The MD simulations also revealed a – to our knowledge – previously unknown binding mode in unsymmetric cyanine dyes related to a partially denatured state of the dsDNA (Figure 7B). We observed a partially denatured state where three terminal base pairs melt and their dynamics are increased. This state (*“denatured, unbound”*) was observed in the absence of dye and was thus not induced by dye binding. The dye readily binds to this highly dynamic region using a combination of electrostatic and π-π interactions with accessible nucleobases (*“denatured, bound” A, B, C, &D*). The MD simulation also revealed the formation of interactions between the dye and a flipped nucleobase (*“flipped”*). These more dynamic regions appear to accommodate a binding mode best described as “dynamic encapsulation”. These observations are intriguing as they offer an alternative explanation as to how these dyes interact with dsDNA and dsRNA, by utilizing localized, partially denatured regions in otherwise double-stranded nucleic acids. These regions are more dynamic than the rigid dsDNA, but not as dynamic as ssDNA. Notably, these observations align with observations reported previously for differential scanning fluorimetry profiles of RNA in the presence of RG (Figure 7A).[33] When the fluorescence of RG in the presence of ssRNA was monitored with rising temperature, the fluorescence decreases and adopts a nearly exponential profile. However, while the fluorescence of RG in the presence of dsRNA initially decreases with rising temperature in a similar fashion, it substantially increases well before reaching the melting temperature of the duplex RNA, around 40-65°C for the example shown in Figure 7A. The current MD simulation data supports the hypothesis that with rising temperature – but well before reaching the melting temperature – more binding sites are created by increased dynamics, e.g. through base flipping and localized denaturation in regions commonly referred to as nucleation bubbles. Fluorescence then sharply returns to the baseline at the melting temperature of the duplex.

**Figure 7:**
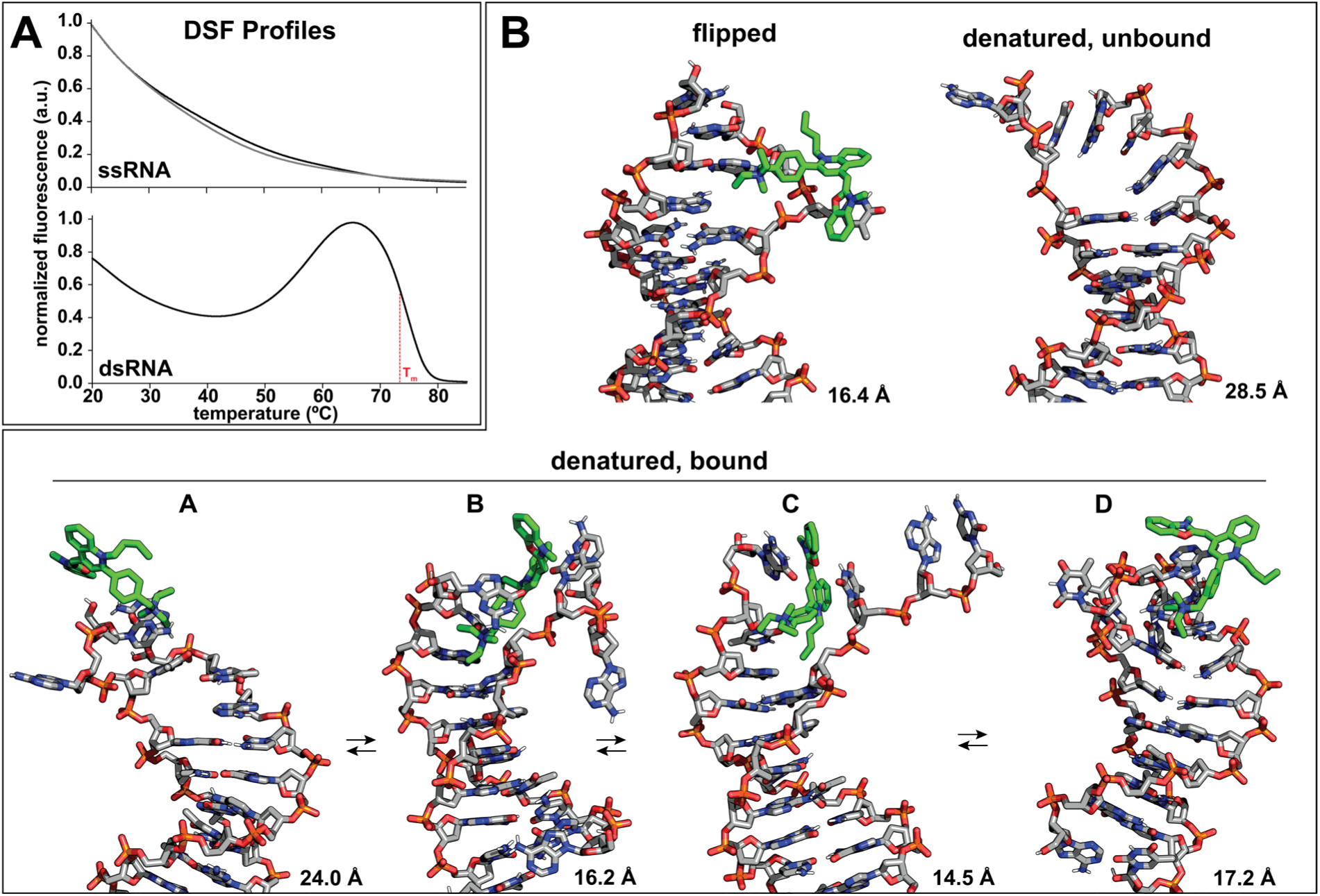
RG binding to denatured DNA states. (A) Differential Scanning Fluorimetry (DSF) profiles of (top) ssRNA and (bottom) dsRNA in the presence of RG. The fluorescence of two disordered ssRNAs (black and gray) shows an approximately exponential decrease with rising temperature, while the dsRNA hybridized from the two ssRNAs displays a substantial rise in fluorescence prior to a sharp decline at the melting temperature (T_m_) of the duplex RNA. Data was reproduced from Silvers et al. [33]. (B) Interactions of RG observed in MD simulations showed interactions of the dye with flipped nucleobases as well as nucleobases present in the partially denatured state of the dsDNA. For each state, the distance between the centers of mass of dye and DNA is shown.

## CONCLUSIONS

The data provided in this study close an important gap in the mechanistic understanding of three widely used commercial unsymmetric cyanine dyes, RG, OG, and PG. By employing high-field NMR spectroscopy and mass spectrometry, we *de novo* established the molecular structures of RG and OG, while independently confirming the previously proposed molecular structure of PG. In doing so, we showed that RG, OG, and PG conform to the general molecular architecture of unsymmetric cyanine dyes that were previously established in other dyes such as TO, SSA, SGO, SGI, and SGII, yet the distinct substituent patterns unique to RG, OG, and PG are likely to influence their binding behavior and photophysical behavior.

Beyond structure determination, this work provides a broader comparative framework for NMR spectroscopic studies of these family of dyes. The complete ^1^H and ^13^C resonance assignments for RG, OG, and PG presented here expand the reference chemical shift set for unsymmetric cyanine dyes. These data sets, together with previously published chemical shift data for five other unsymmetric cyanine dyes (TO, SSA, SGO, SGI, and SGII) should facilitate future structural studies of related commercial and non-commercial fluorophores alike.

The photophysical data further demonstrate that all three dyes behave as strongly fluorogenic probes whose signal increases markedly upon binding nucleic acids, with the greatest enhancement observed for double-stranded substrates. These findings help rationalize their analytical utility while also clarifying that differences in marketed applications do not necessarily reflect simple, exclusive selectivity for a single nucleic acid class or type.

Importantly, NMR spectroscopic data and long-timescale molecular dynamics simulations provided an in-depth mechanistic picture of dominant interactions between dye and double-stranded DNA. Overall, the interaction between the dyes and the nucleic acid is dominated by a highly dynamic sampling of a variety of binding modes and locations rather than by a single uniform binding mode or location. These results support a model in which binding is governed primarily by a combination of electrostatic interactions with the phosphodiester backbone as well as π-π stacking with accessible nucleobases of the DNA duplex. Besides binding to terminal base pairs of the dsDNA, MD simulations suggest that accessibility to nucleobases is created by dynamics, such as opening of the terminal base pair, base flipping, as well as localized denaturation creating binding sites. This is also significant in the broader context of nucleic acid binding because it highlights that the performance of these dyes depends not only on their molecular structures, but also on nucleic acid topology and local conformational dynamics on multiple timescales.

Overall, the work presented here establishes a molecular foundation for more informed use of RG, OG, and PG in biochemical, biophysical, and analytical applications. It also contributes to the larger goal of connecting fluorophore structure to binding mechanisms and fluorescence response, an essential step toward the rational selection and design of nucleic acid dyes for specific experimental settings.

## Supporting information

MD simulation (Run1)

MD simulation (Run2)

## DATA AVAILABILITY

The ^1^H, ^13^C resonance assignments of the dyes RG, OG, and PG are provided in the Supplementary Information and the molecular dynamics simulations data is provided as Supplementary Files.

## SUPPLEMENTARY DATA

Supplementary data is available at NAR online.

## ACKNOWLEDGEMENTS

Research reported in this publication was supported by NIGMS of the National Institutes of Health under award number R35GM142912 (R.S.). The content is solely the responsibility of the authors and does not necessarily represent the official views of the National Institutes of Health. W.Y. acknowledges funding support from the National Institutes of Health (R01GM124621 and R01GM147673). We thank Dr. Peter Randolph (FSU) for his assistance with absorbance and fluorescence measurements.

## AUTHOR CONTRIBUTIONS

**Nolan Blackford**: Investigation, Methodology, Formal analysis, Data Curation, Visualization, Writing – original draft, Writing – review & editing. **Saileena Nepal**: Investigation, Formal analysis, Data curation, Writing – original draft, Writing – review & editing. **Huan He**: Investigation, Formal analysis, Writing – review & editing. **Lianqing Zheng**: Investigation, Formal analysis, Writing – review & editing. **Wei Yang**: Investigation, Formal analysis, Writing – review & editing. **Robert Silvers**: Conceptualization, Data Curation, Formal analysis, Funding acquisition, Investigation, Methodology, Project administration, Supervision, Validation, Visualization, Writing – original draft, Writing – review & editing.

## Abbreviations

RG: RiboGreen
OG: OliGreen
PG: PicoGreen
SGI: SYBR Green I
SGII: SYBR Green II
SGO: SYBR Gold
SSA: SYBR Safe
TO: Thiazole Orange
YO: Oxazole Yellow

