## Supporting Information for "Molecular Structure and DNA Binding Mode of Unsymmetric Cyanine Dyes RiboGreen and OliGreen"

| | |
|---|---|
| Nolan Blackford | 0009-0008-5818-3870 |
| Saileena Nepal | 0009-0004-0591-0393 |
| Huan He | 0000-0002-5014-9124 |
| Lianqing Zheng | 0009-0000-9077-6889 |
| Wei Yang | 0000-0002-4520-3253 |
| Robert Silvers | 0000-0003-0197-3878 |

### Supporting Figures

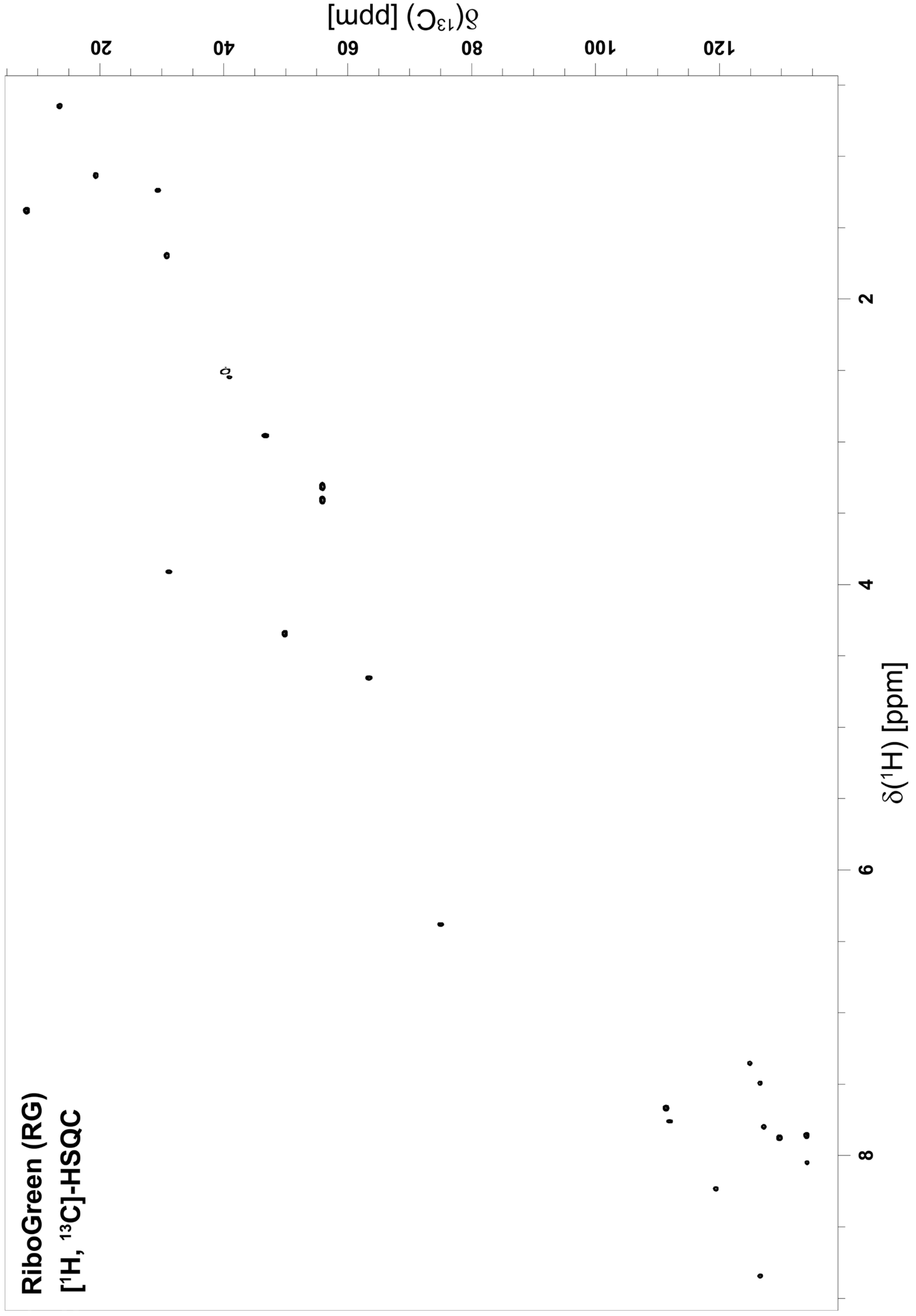


**Figure S1:** [$^1$H, $^{13}$C]-HSQC of RiboGreen recorded in DMSO-$d_6$ at 700 MHz.

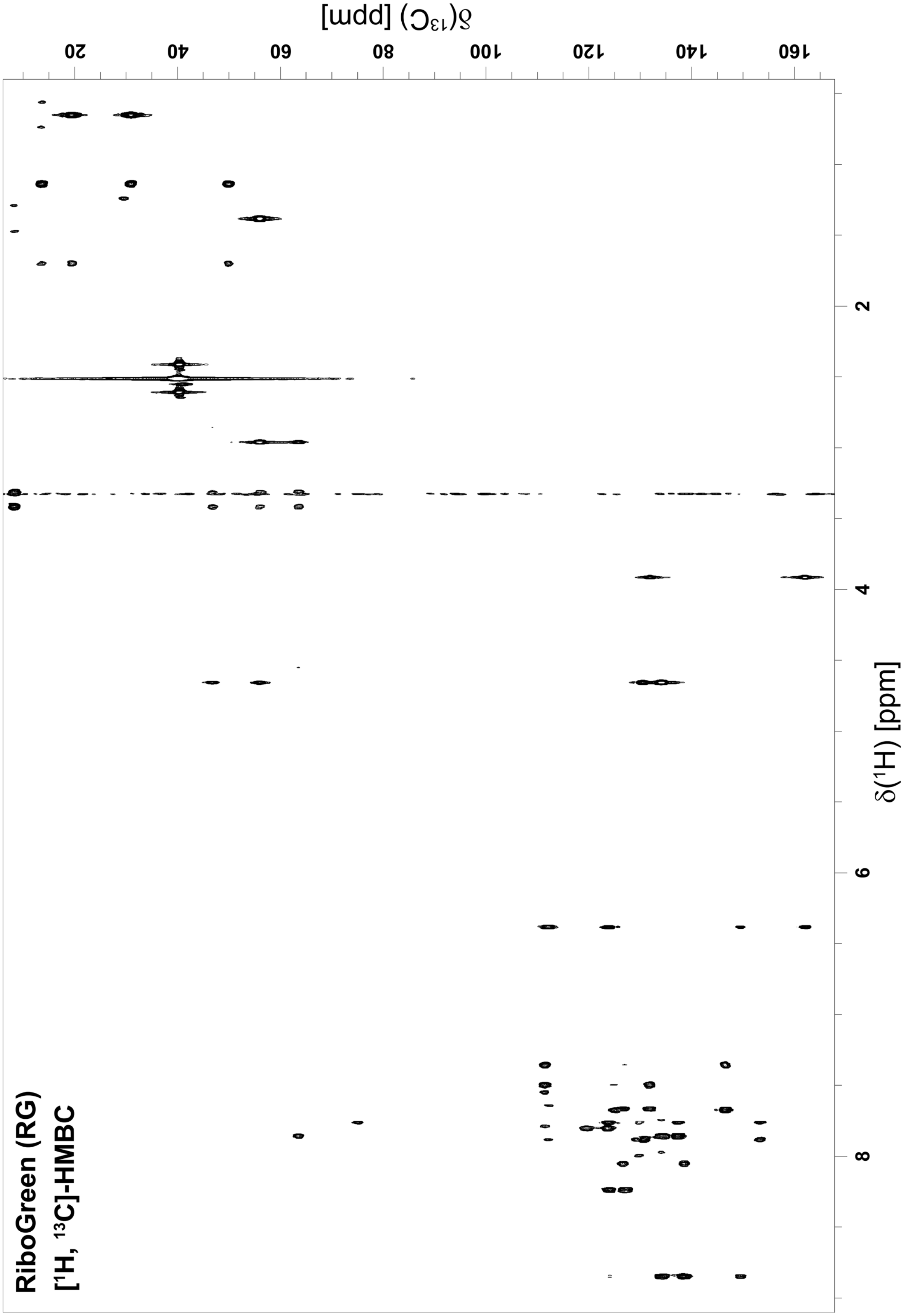


**Figure S2:** [$^{1}$H, $^{13}$C]-HMBC of RiboGreen recorded in DMSO-$d_6$ at 700 MHz.

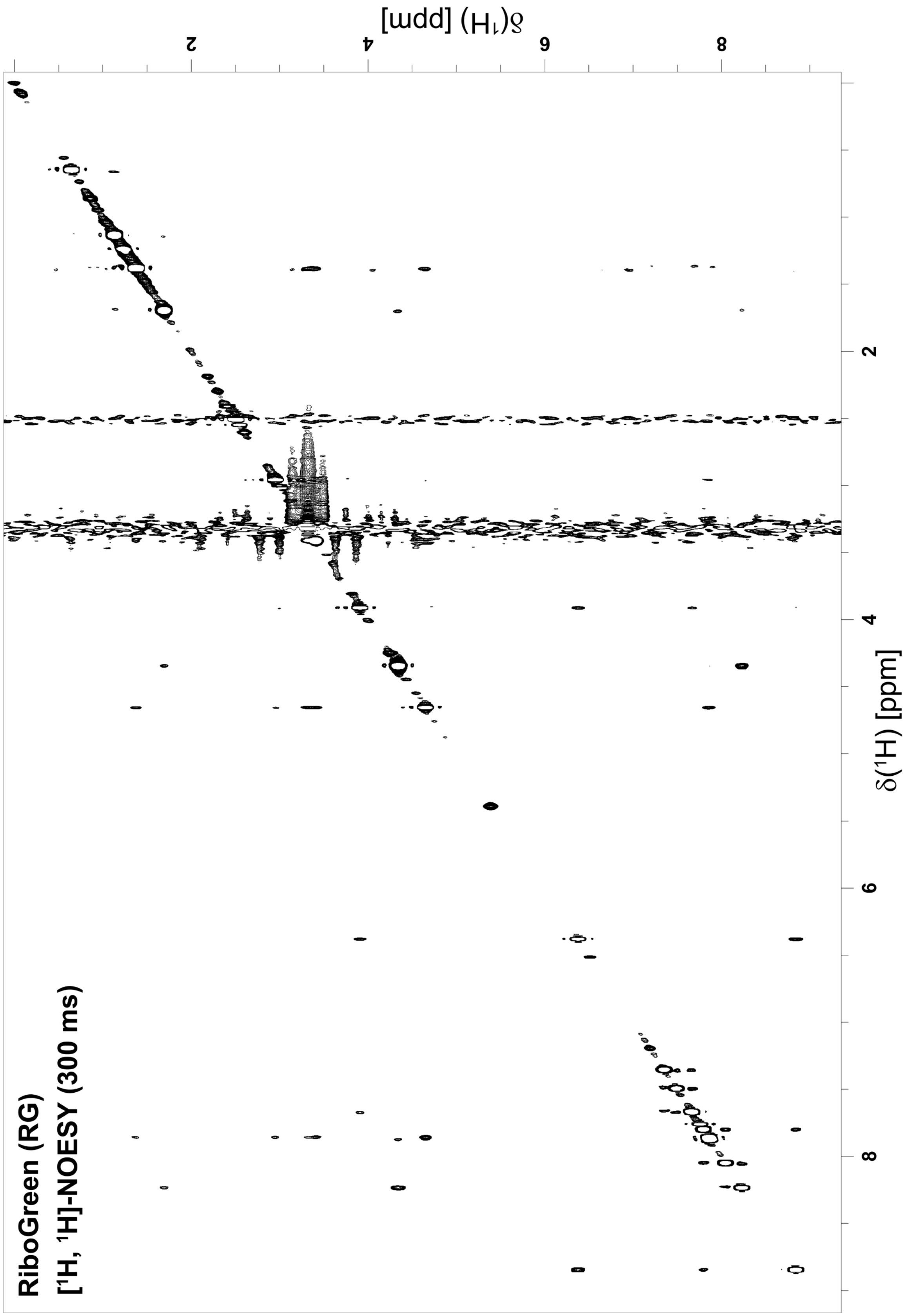


**Figure S3:** [$^1$H, $^1$H]-NOESY of RiboGreen recorded in DMSO-$d_6$ at 700 MHz.

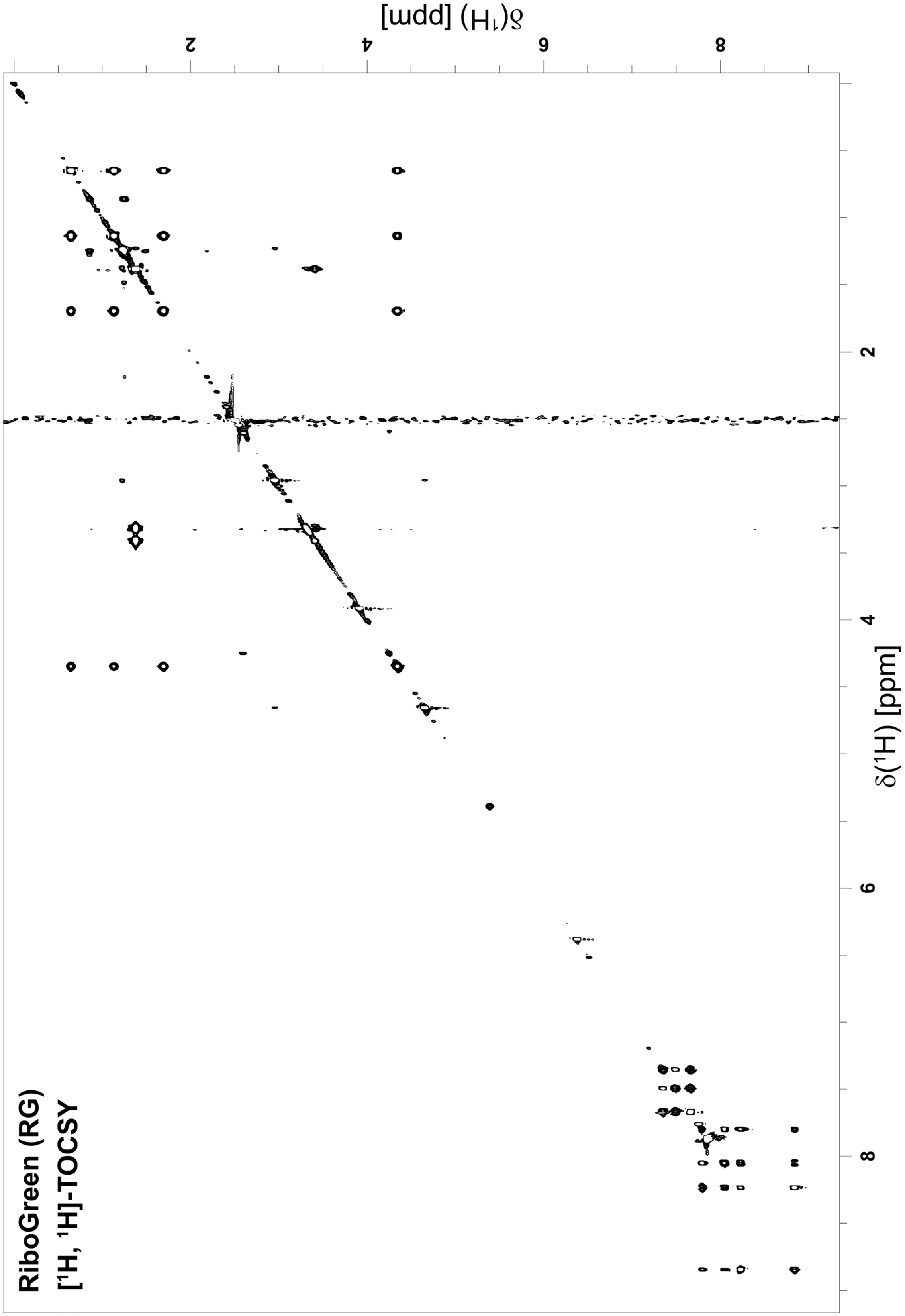


**Figure S4:** [$^1$H, $^1$H]-TOCSY of RiboGreen recorded in DMSO-d$_6$ at 700 MHz.

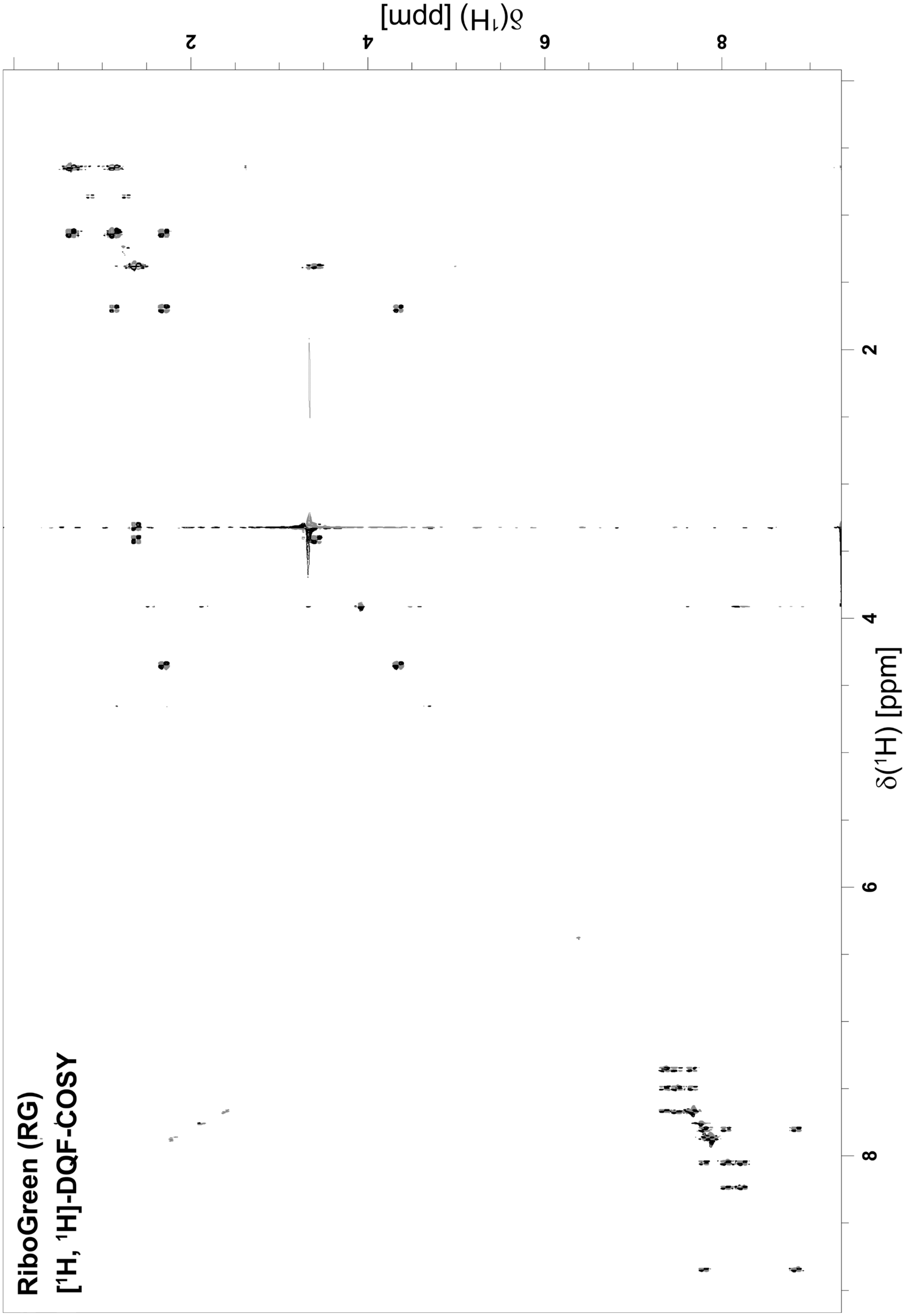


**Figure S5:** [$^{1}$H, $^{1}$H]-DQF-COSY of RiboGreen recorded in DMSO-d$_6$ at 700 MHz. Positive peaks are black and negative peaks are gray.

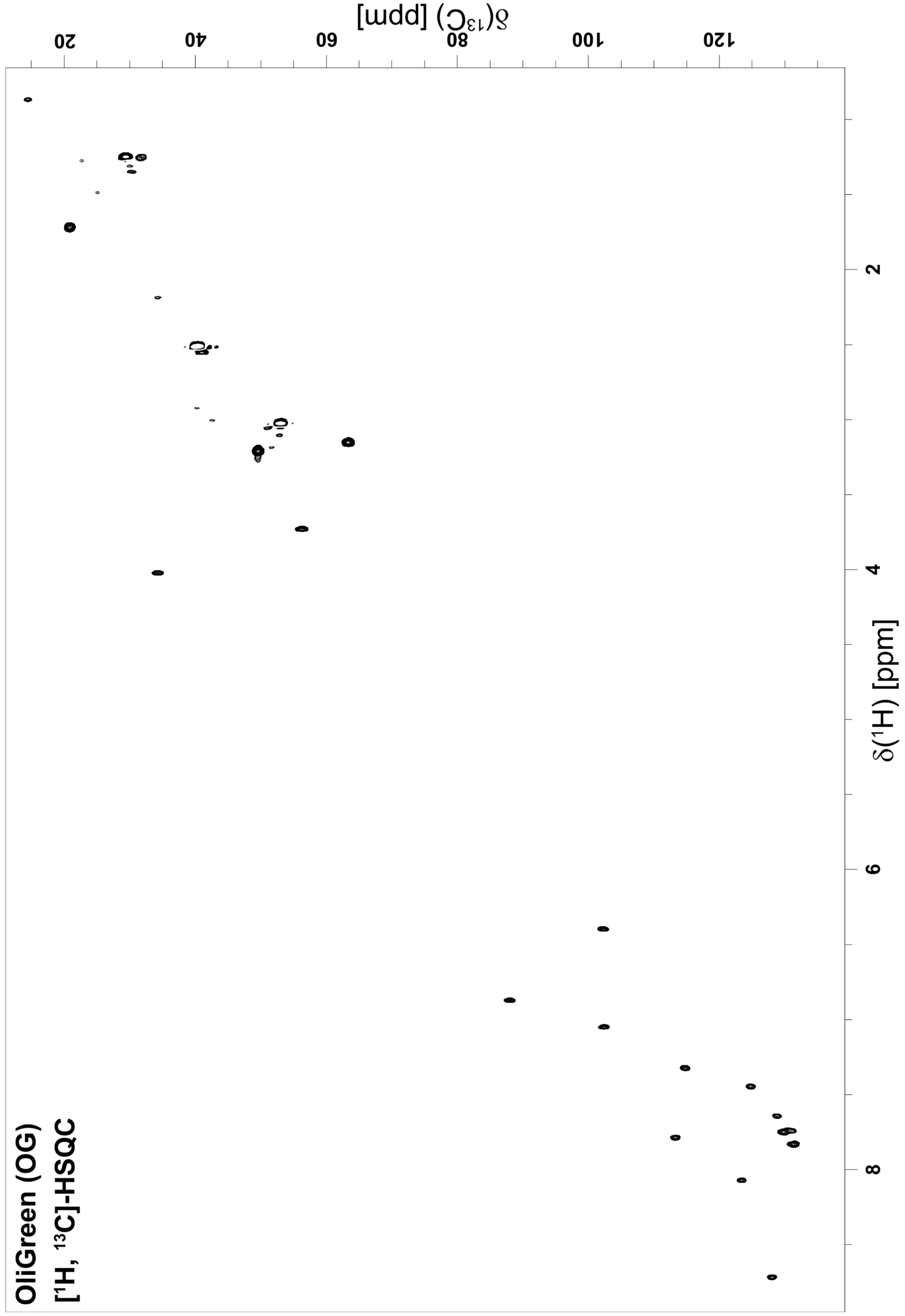


**Figure S6:** [$^1$H, $^{13}$C]-HSQC of OliGreen recorded in DMSO-$d_6$ at 700 MHz.

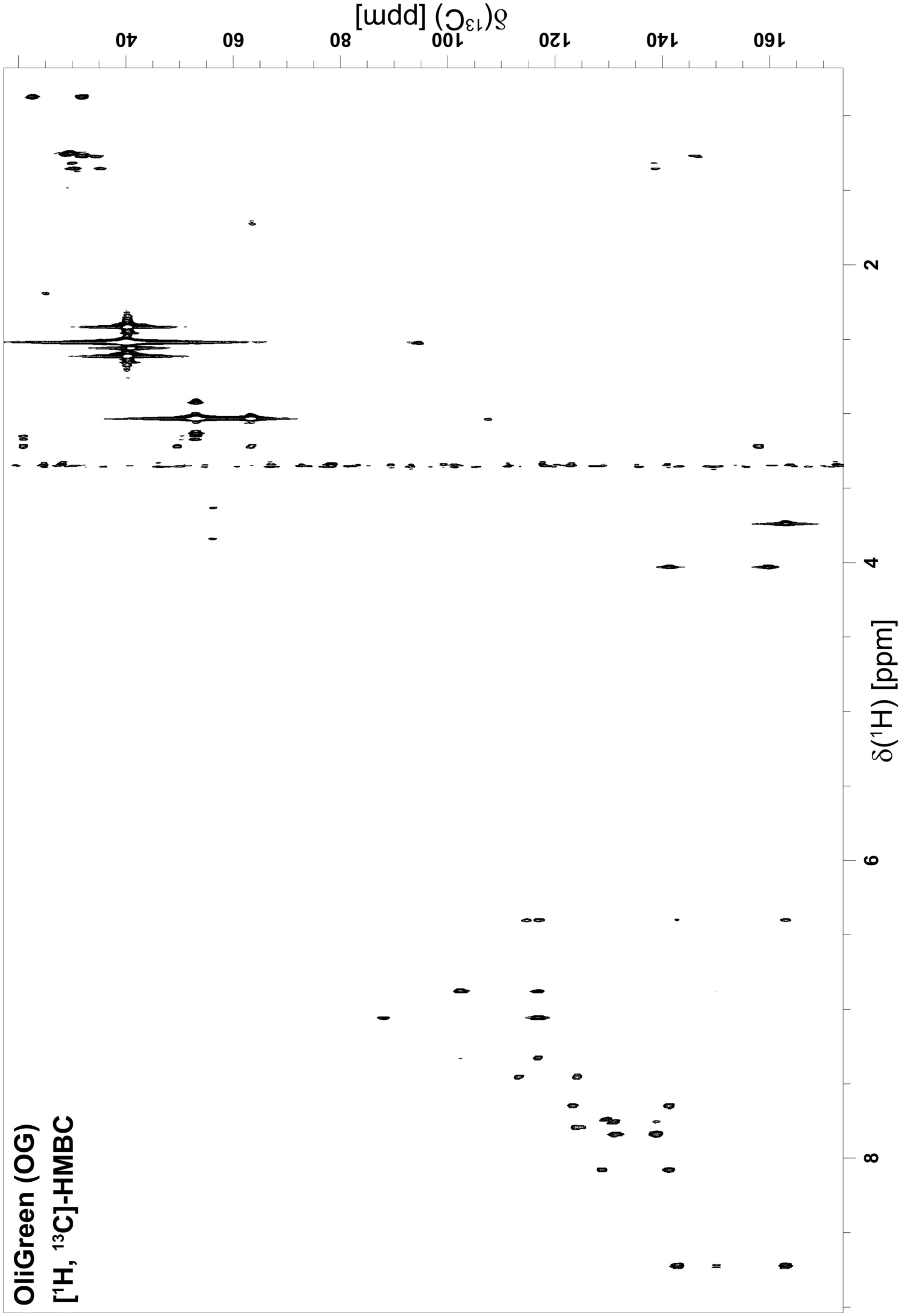


**Figure S7:** [$^1$H, $^{13}$C]-HMBC of OliGreen recorded in DMSO-d$_6$ at 700 MHz.

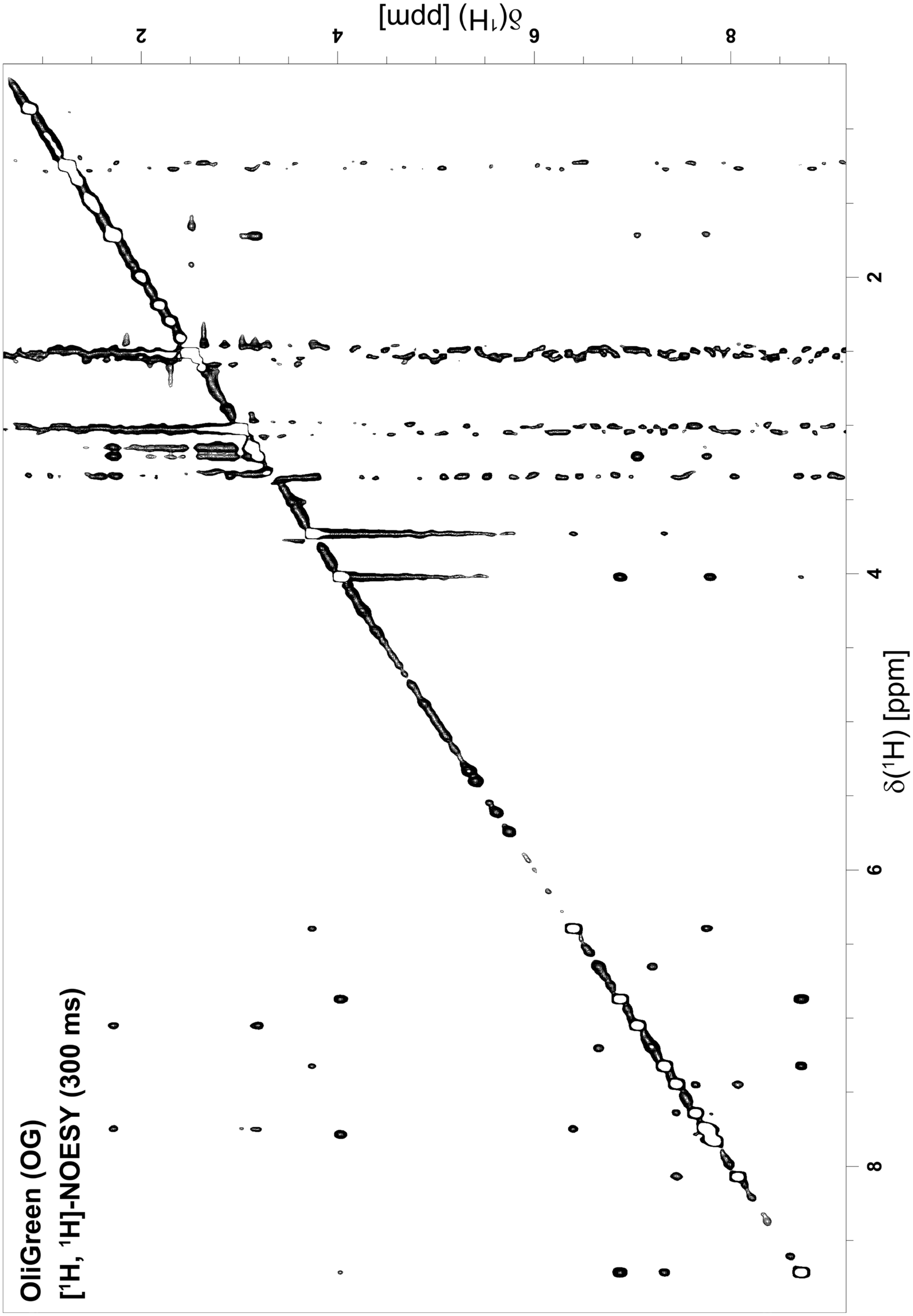


**Figure S8:** [$^1$H, $^1$H]-NOESY of OliGreen recorded in DMSO-d$_6$ at 700 MHz.

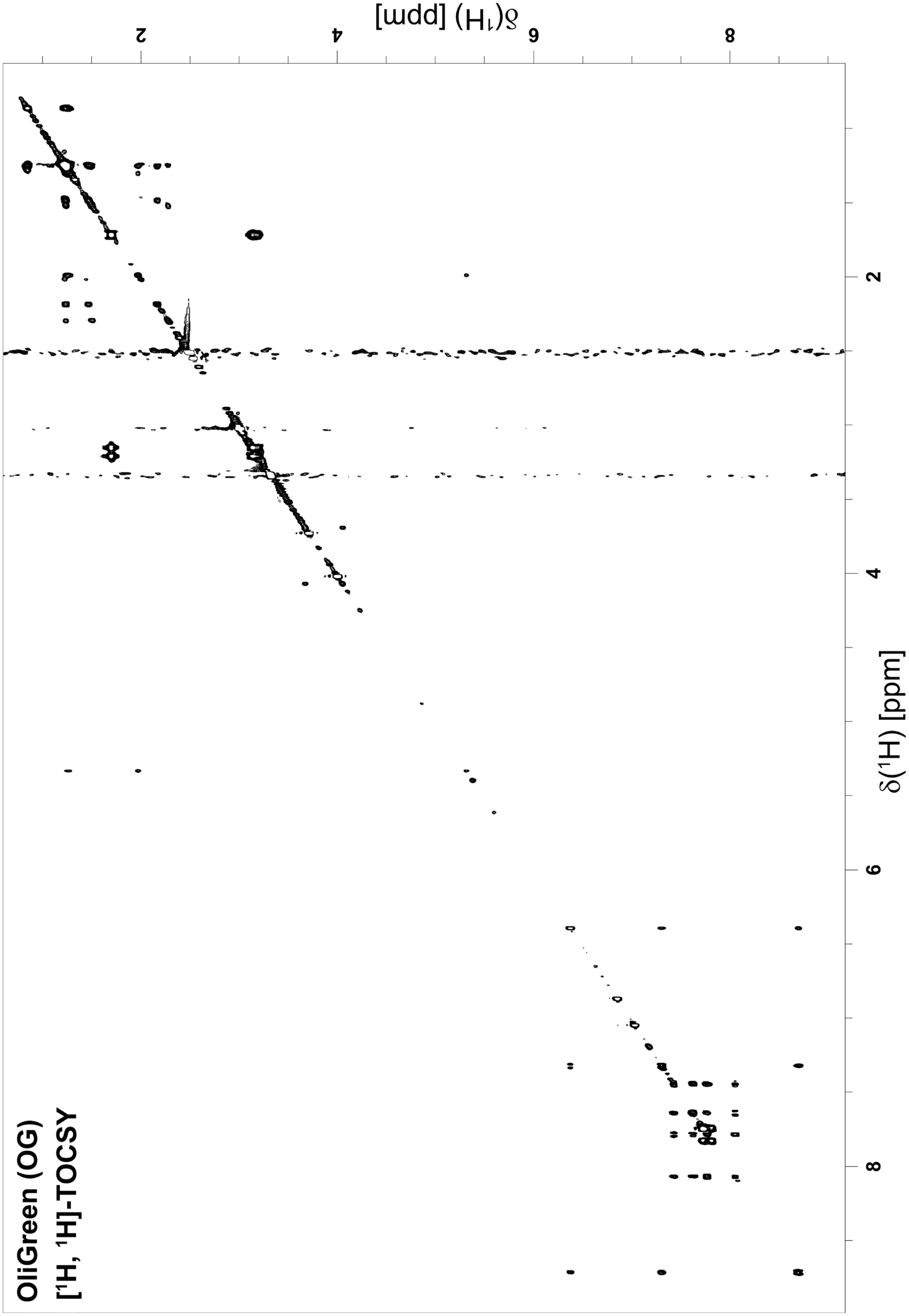


**Figure S9:** [$^{1}$H, $^{1}$H]-TOCSY of OliGreen recorded in DMSO-d$_6$ at 700 MHz.

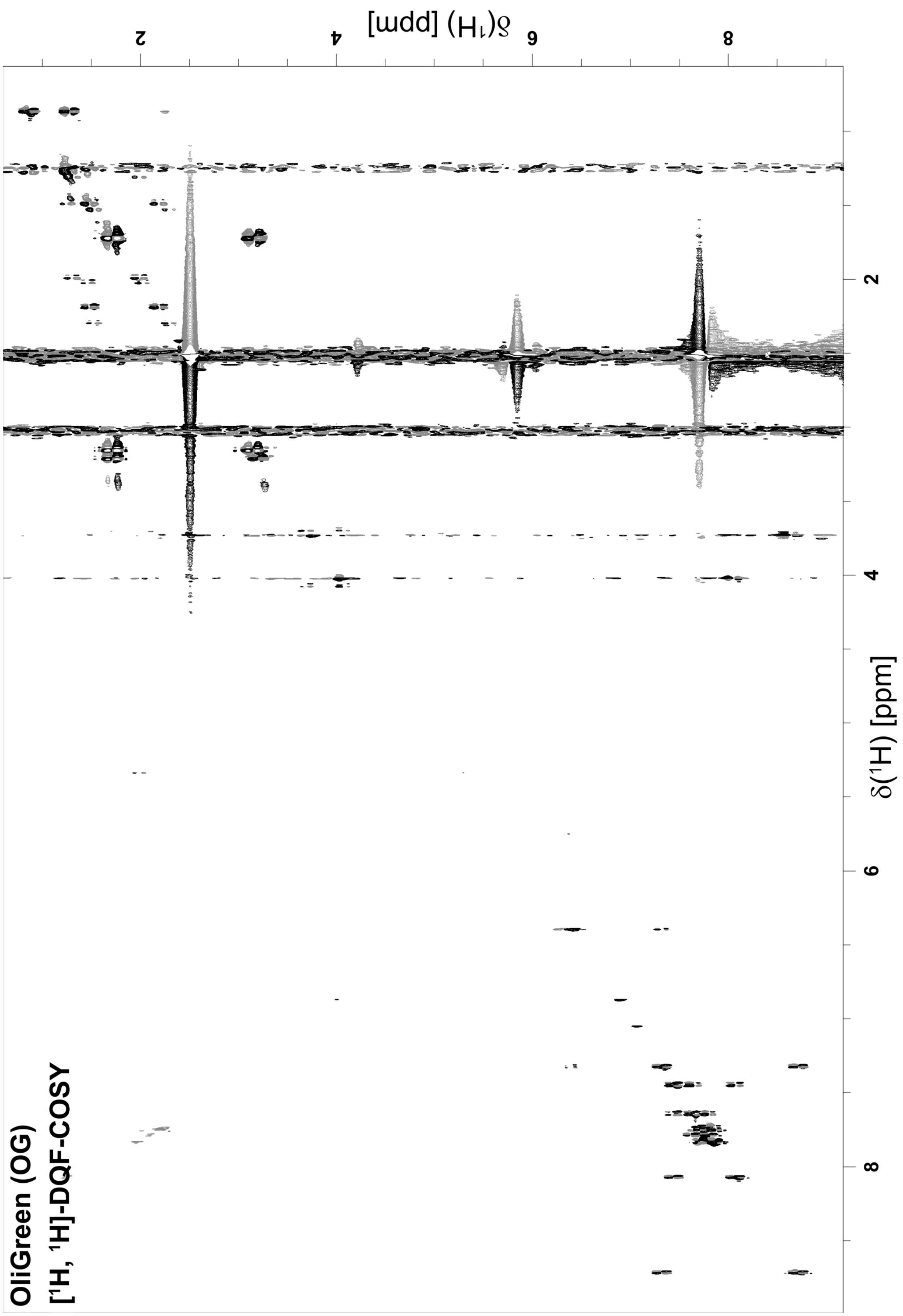


**Figure S10:** [$^1$H, $^1$H]-DQF-COSY of OliGreen recorded in DMSO-$d_6$ at 700 MHz. Positive peaks are black and negative peaks are gray.

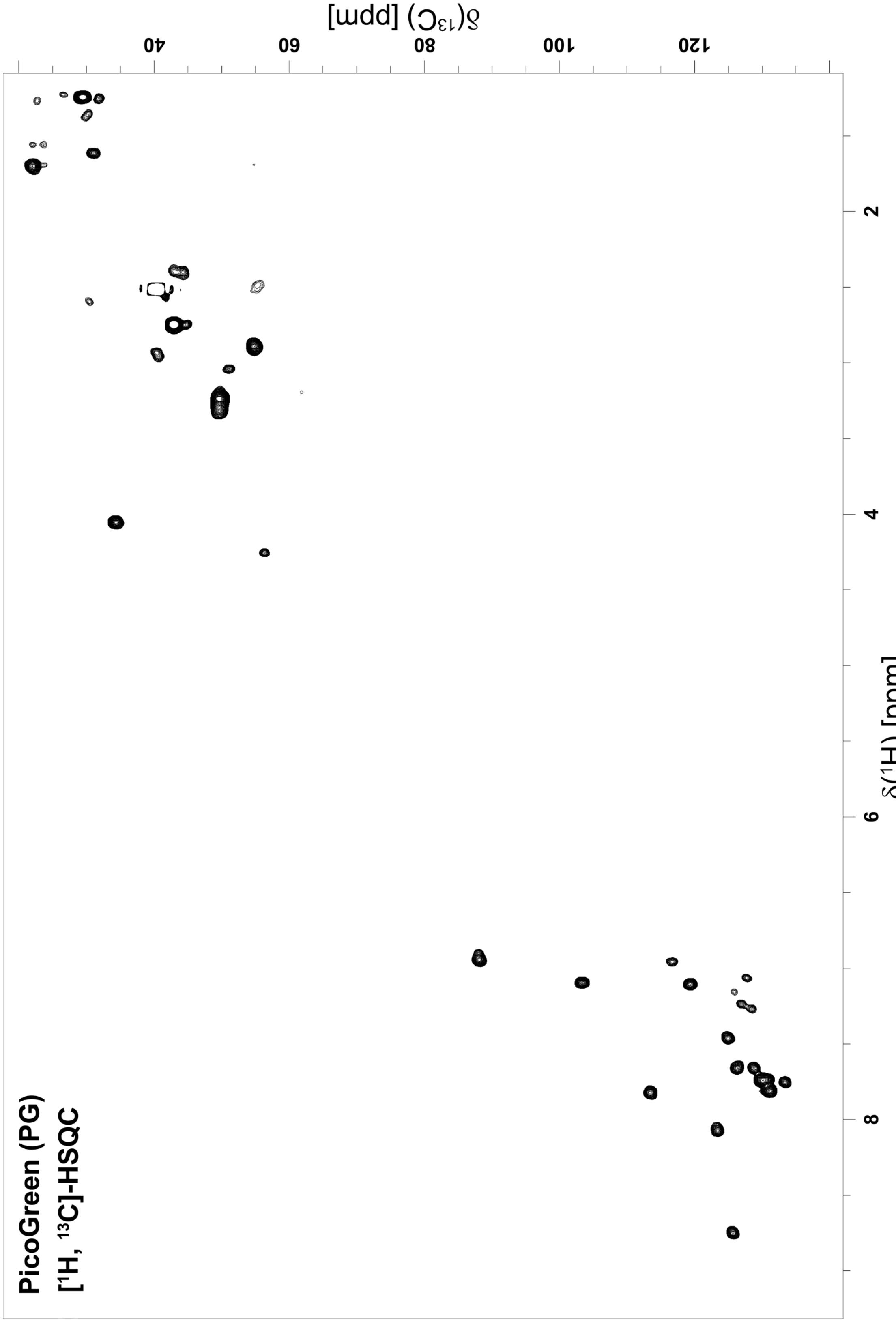


**Figure S11:** [$^1$H, $^{13}$C]-HSQC of PicoGreen recorded in DMSO-$d_6$ at 700 MHz.

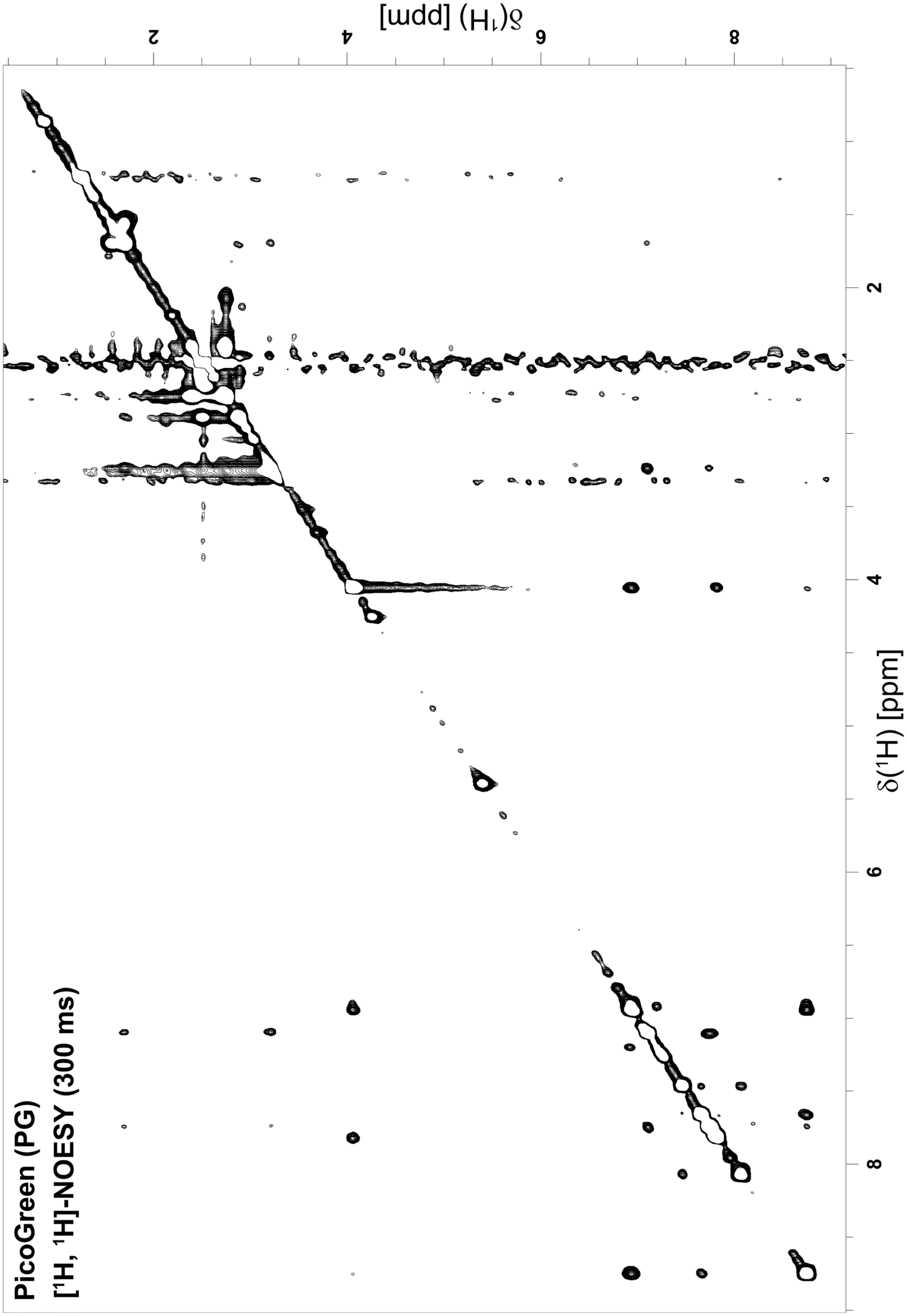


**Figure S12:** [$^1$H, $^1$H]-NOESY of PicoGreen recorded in DMSO-$d_6$ at 700 MHz.

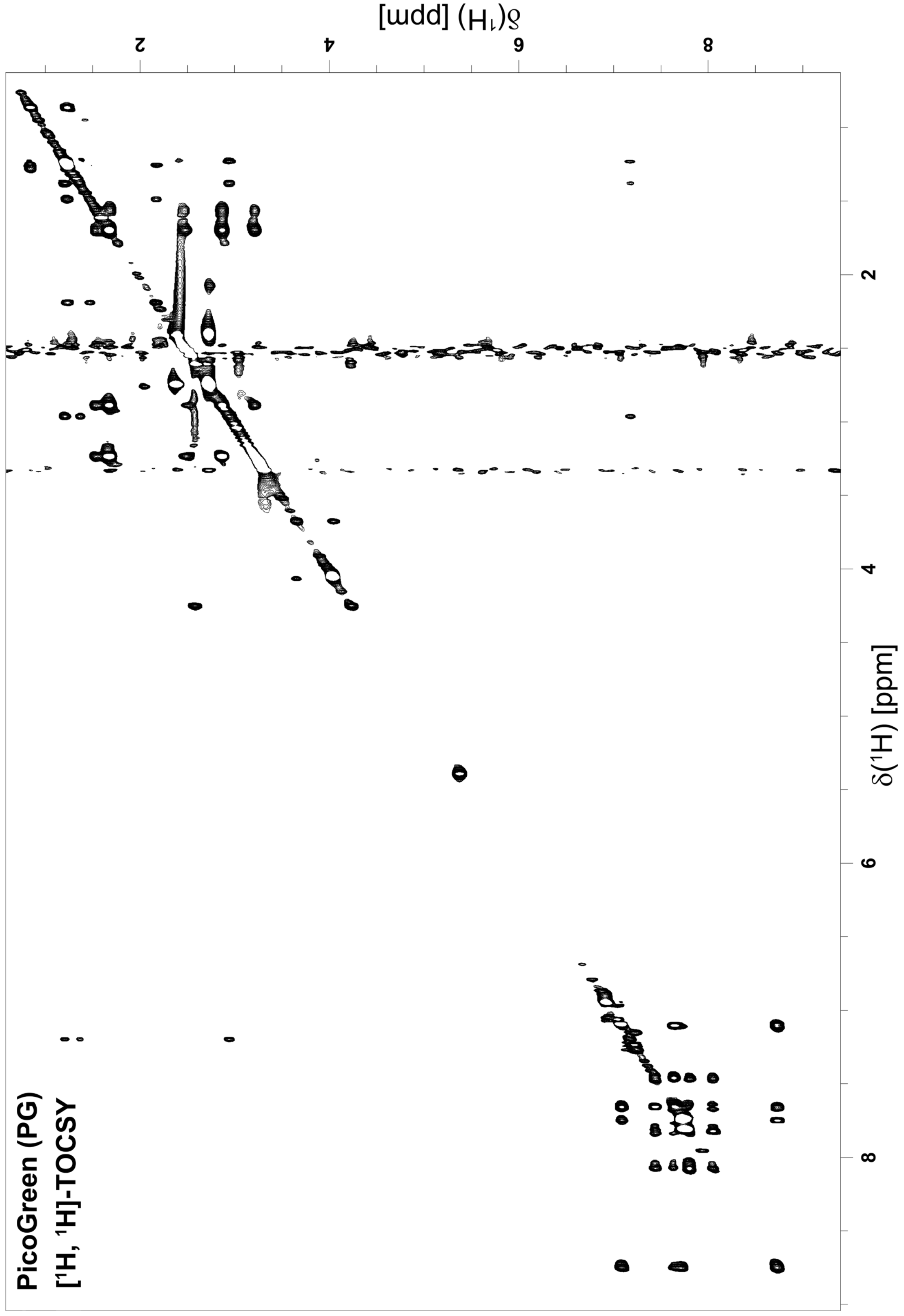


**Figure S13:** [$^1$H, $^1$H]-TOCSY of PicoGreen recorded in DMSO-d$_6$ at 700 MHz.

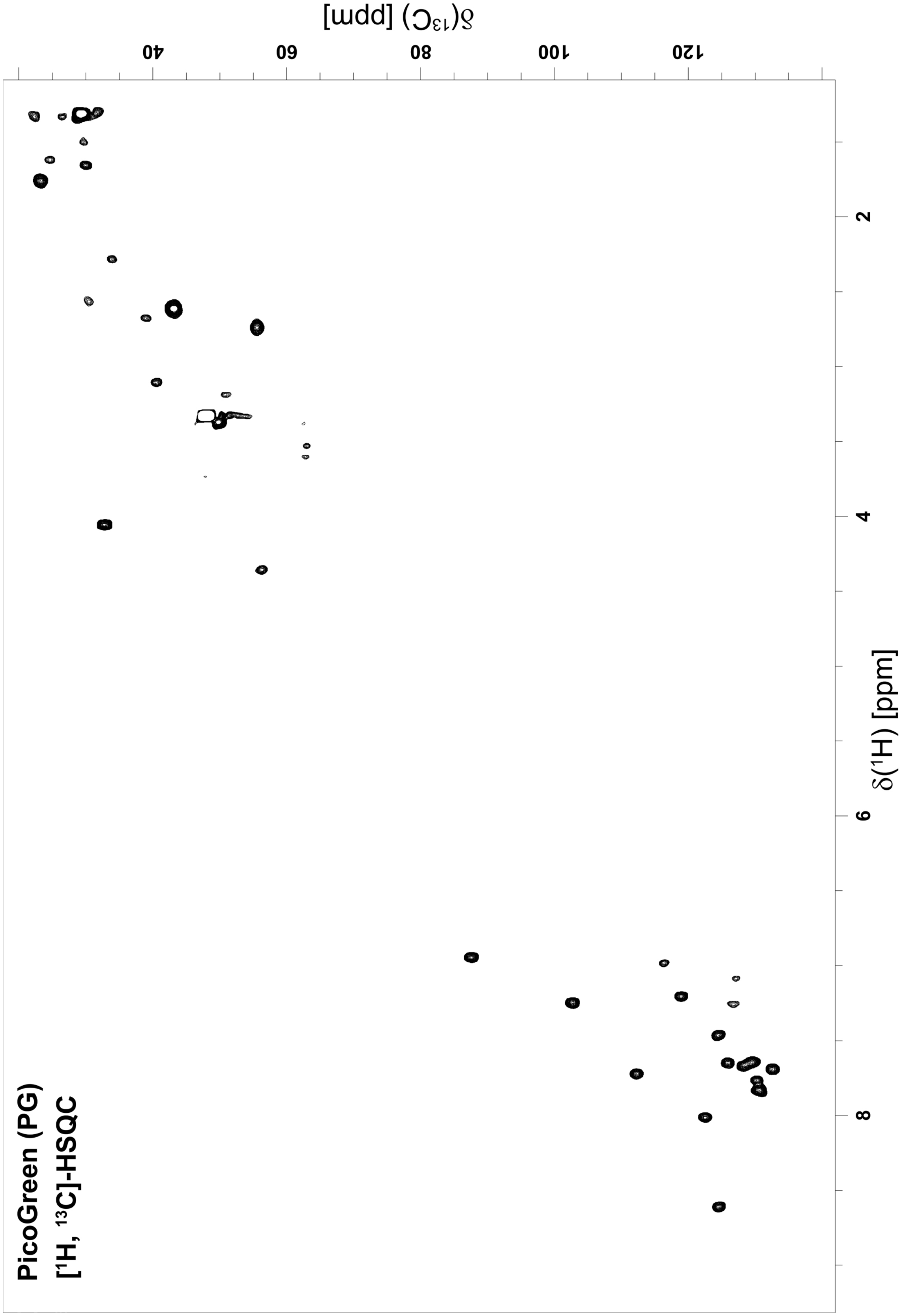


**Figure S14:** [$^1$H, $^{13}$C]-HSQC of PicoGreen recorded in methanol-d$_4$ at 700 MHz.

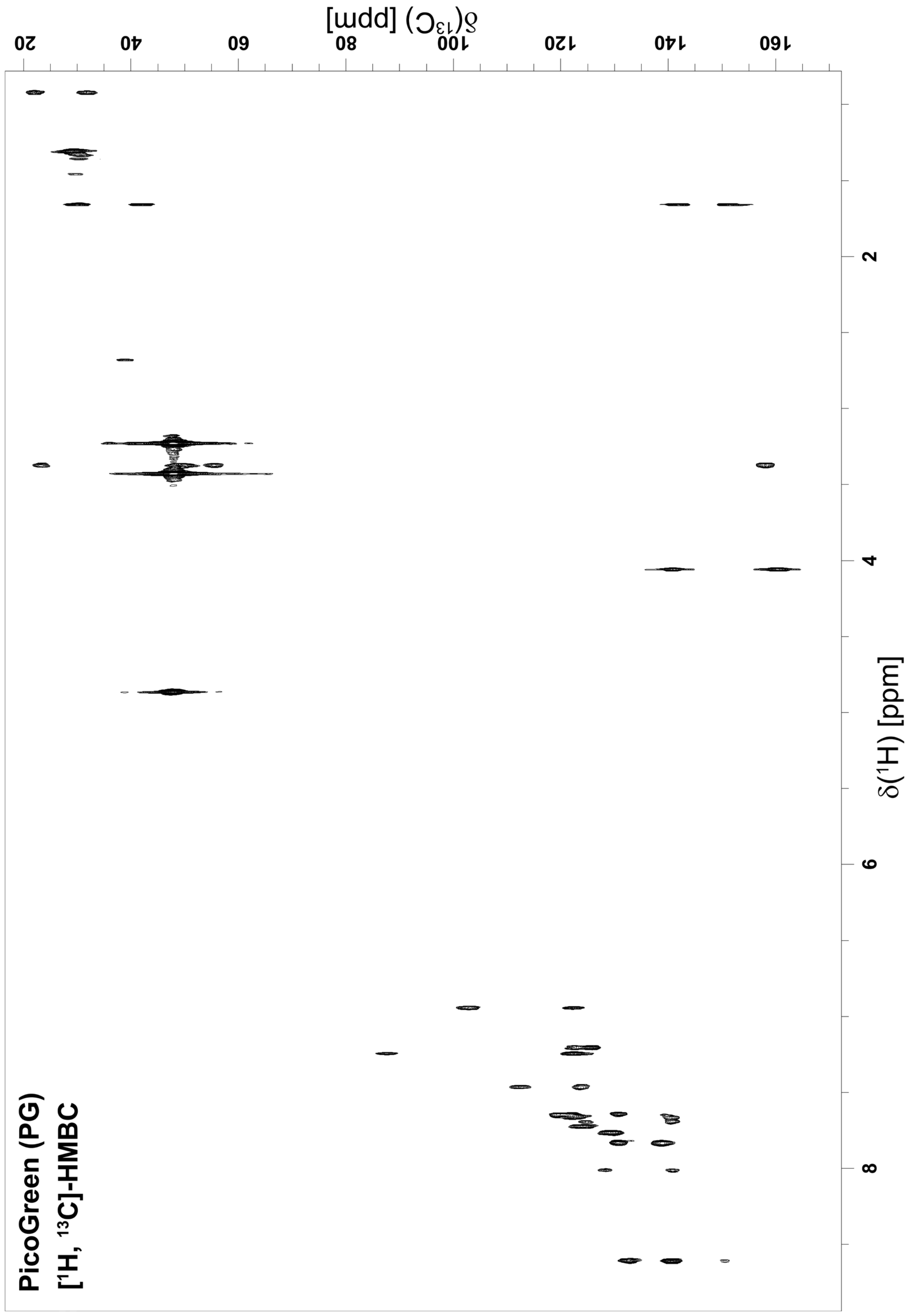


**Figure S15:** [$^{1}$H, $^{13}$C]-HMBC of PicoGreen recorded in methanol-$d_4$ at 700 MHz.

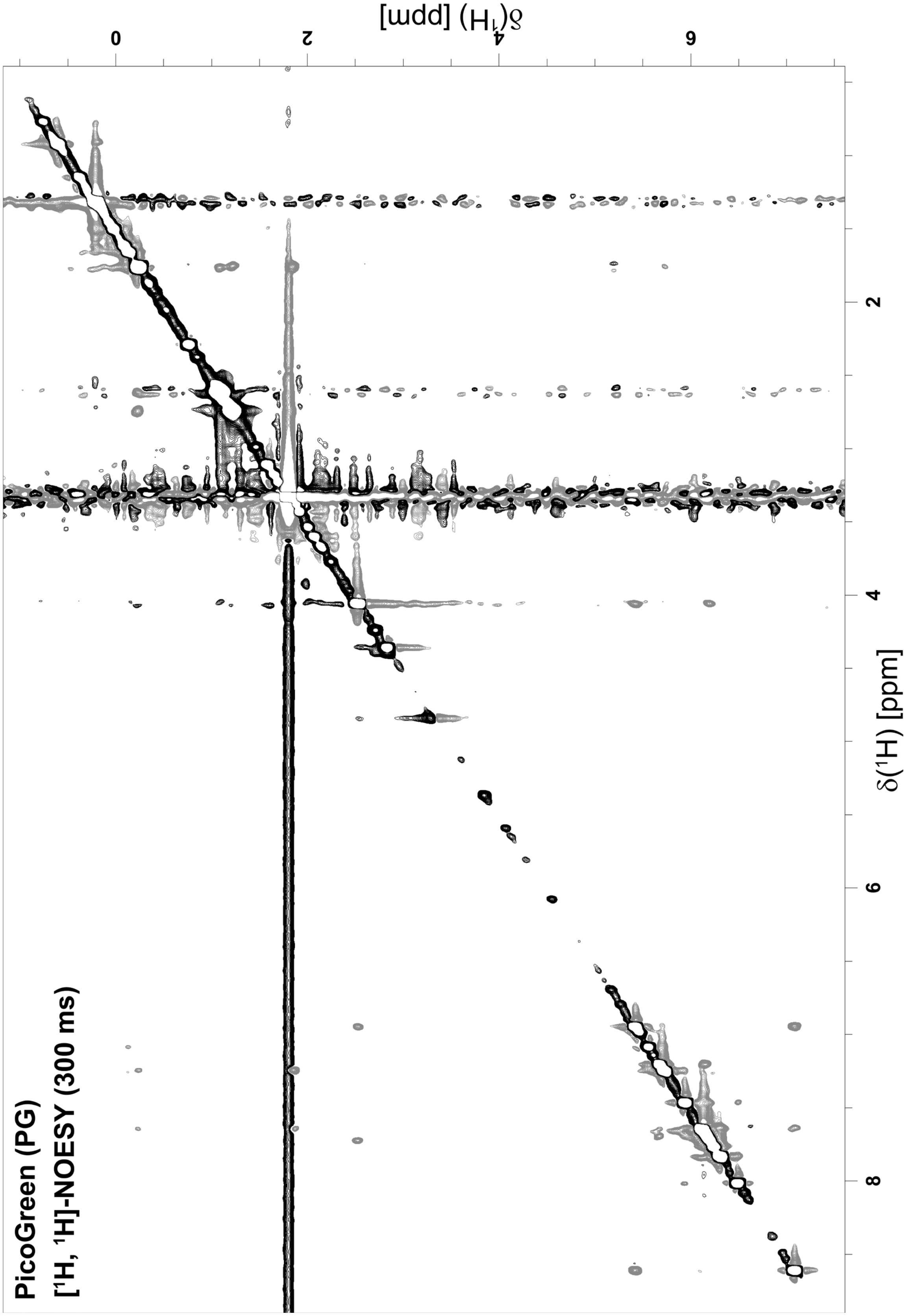


**Figure S16:** [$^{1}$H, $^{1}$H]-NOESY of PicoGreen recorded in methanol-$d_4$ at 700 MHz. Positive peaks are black and negative peaks are gray.

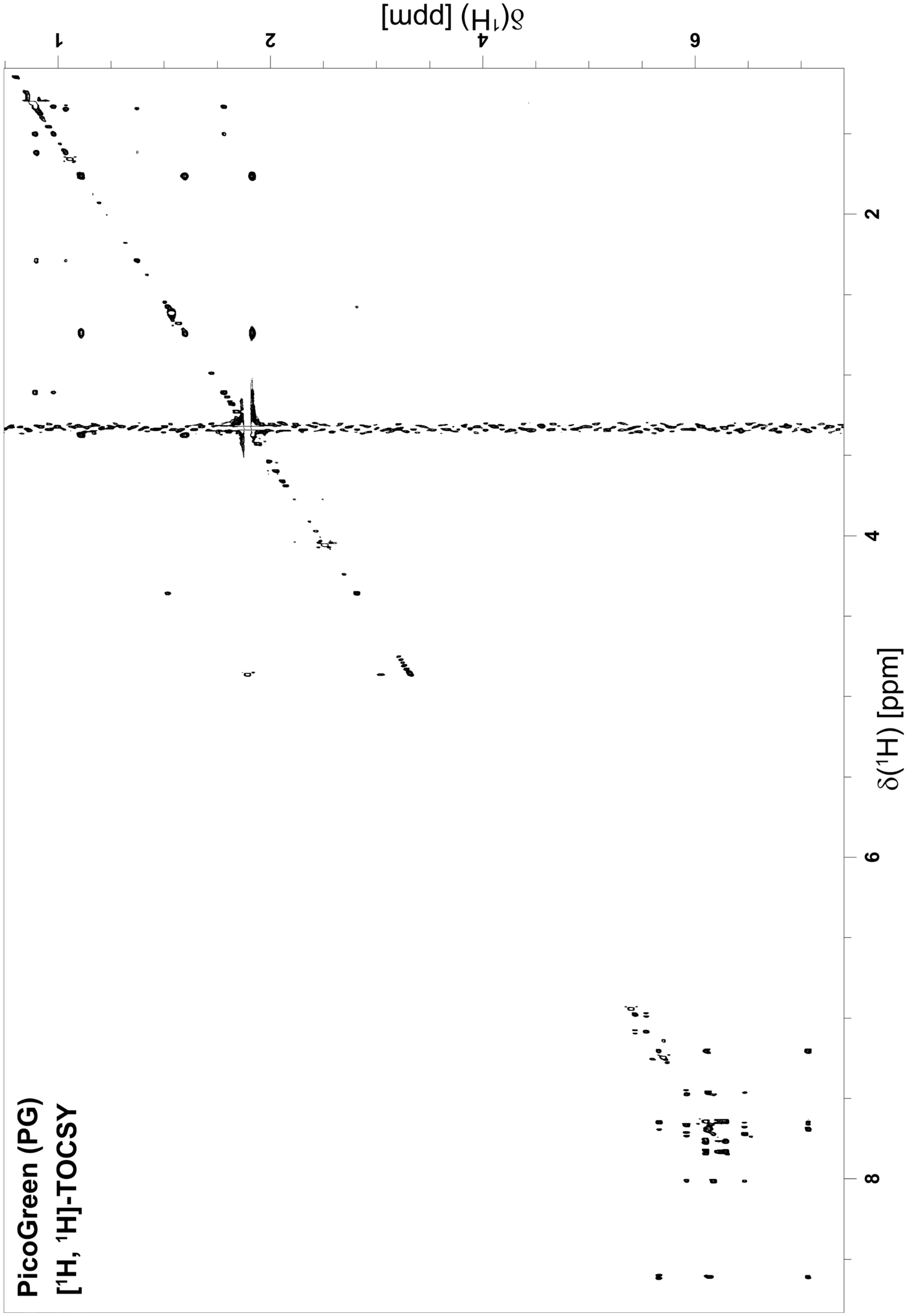


**Figure S17:** [$^1$H, $^1$H]-TOCSY of PicoGreen recorded in methanol-$d_4$ at 700 MHz.

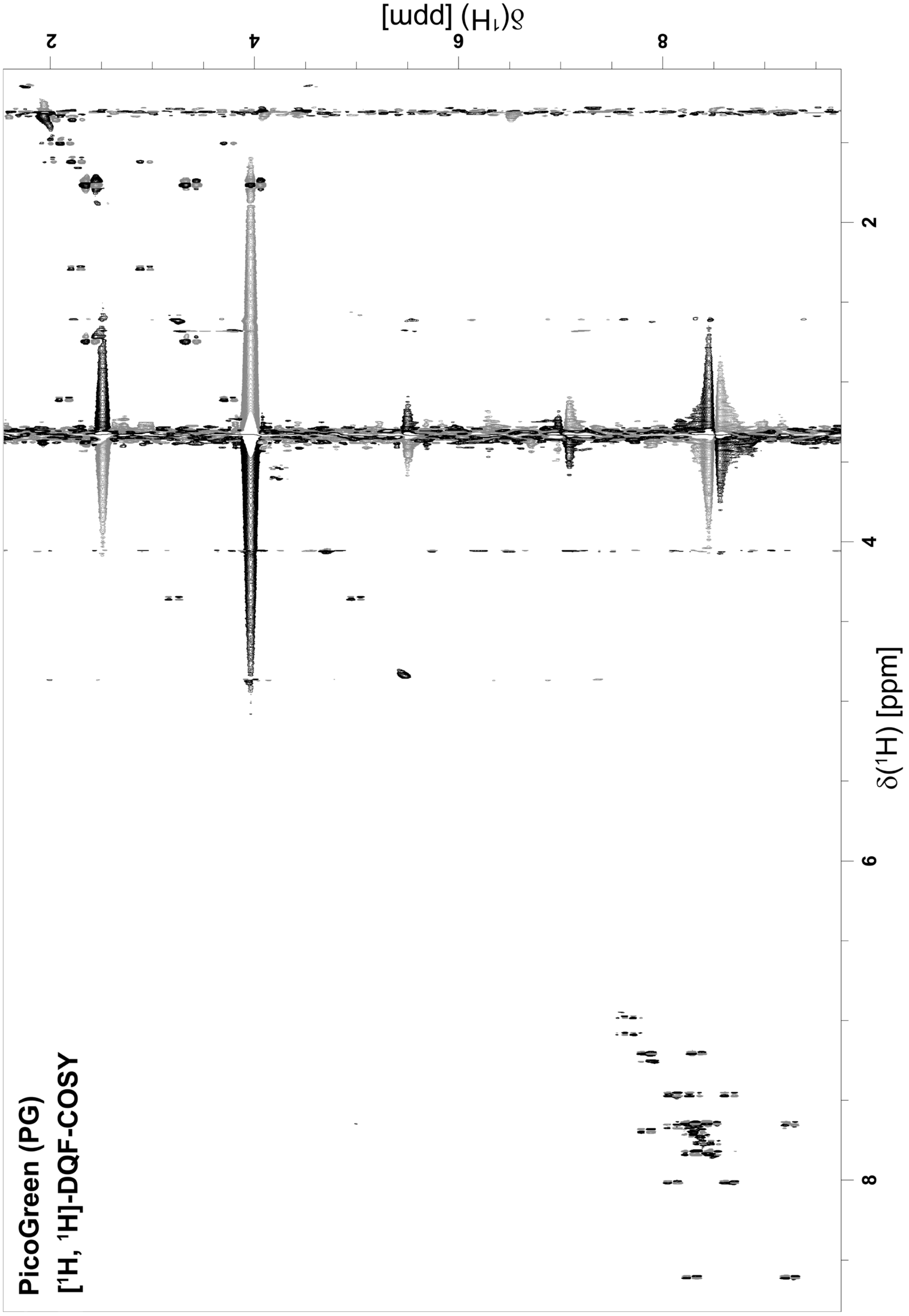


**Figure S18:** [$^1$H, $^1$H]-DQF-COSY of PicoGreen recorded in methanol-d$_4$ at 700 MHz. Positive peaks are black and negative peaks are gray.

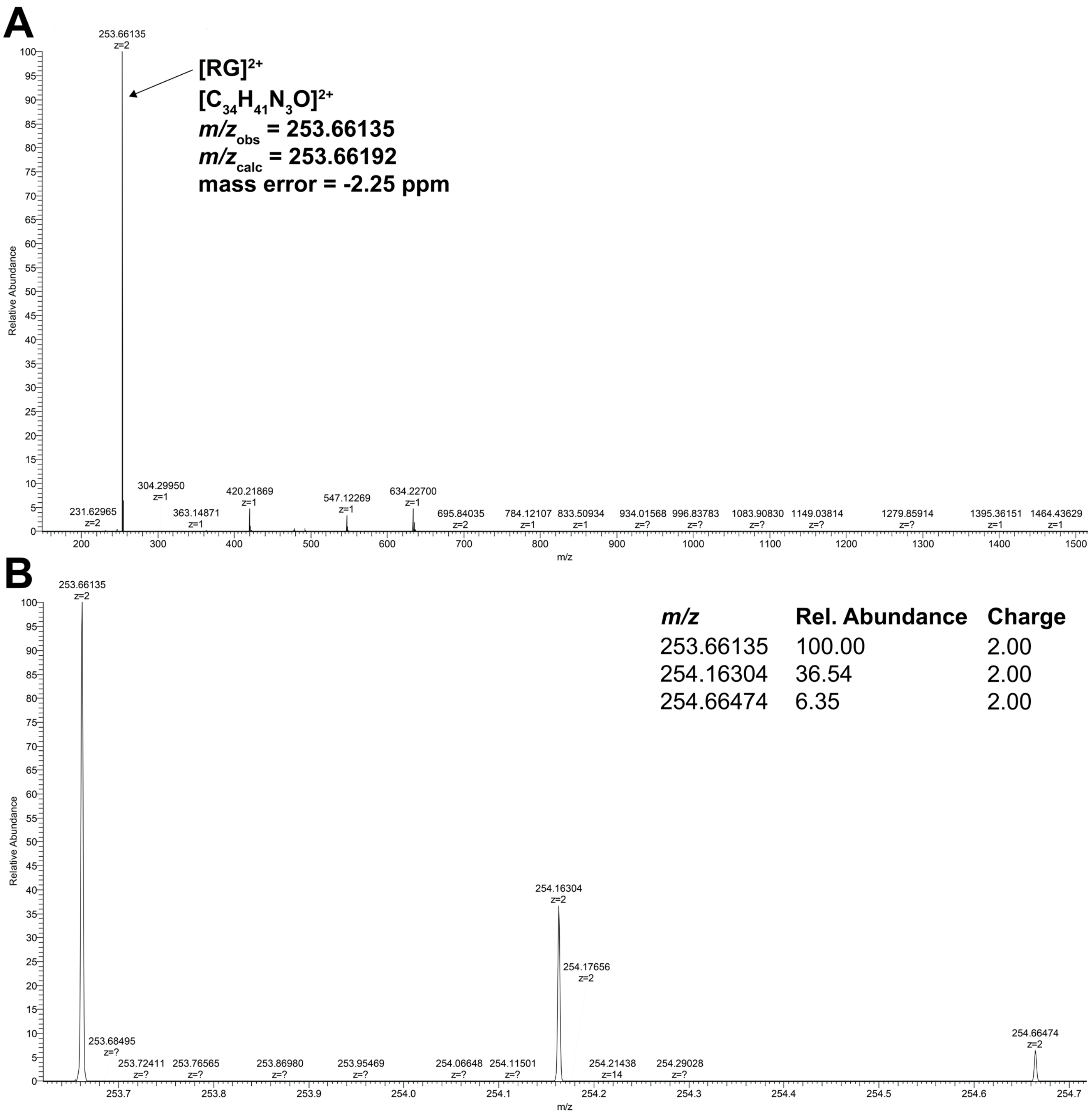


**Figure S19**: MS analysis of RiboGreen (RG). (A) ESI$^+$ mass spectrum with m/z range of 150.0-1500.0. The observed and calculated m/z ratios of the main species [RG]$^{2+}$ are shown. (B) Zoom of the ESI$^+$ mass spectrum(A) showing the isotopic distribution of the main species [RG]$^{2+}$. m/z ratios, relative abundances, and charges of observed peaks are as listed.

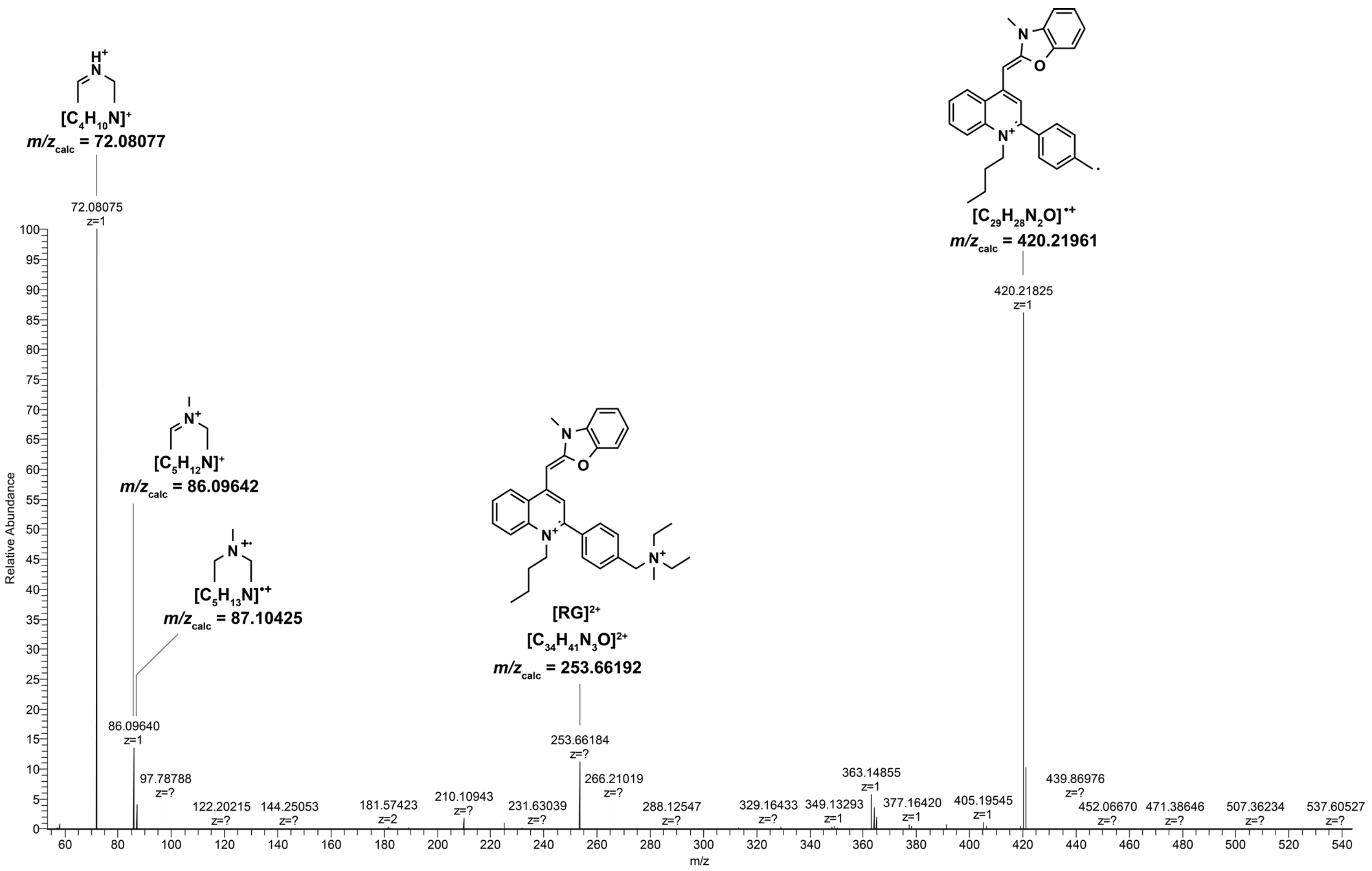


**Figure S20**: MS/MS analysis of RiboGreen (RG). The tandem MS spectrum (253.0 at 40% HCD) with m/z range of 53.816-538.160 shows the presence of several fragments of the precursor species $[RG]^{2+}$. Structures, empirical formulae, and calculated m/z ratios are shown for the precursor peak (m/z 253.66184) as well as select fragment peaks.

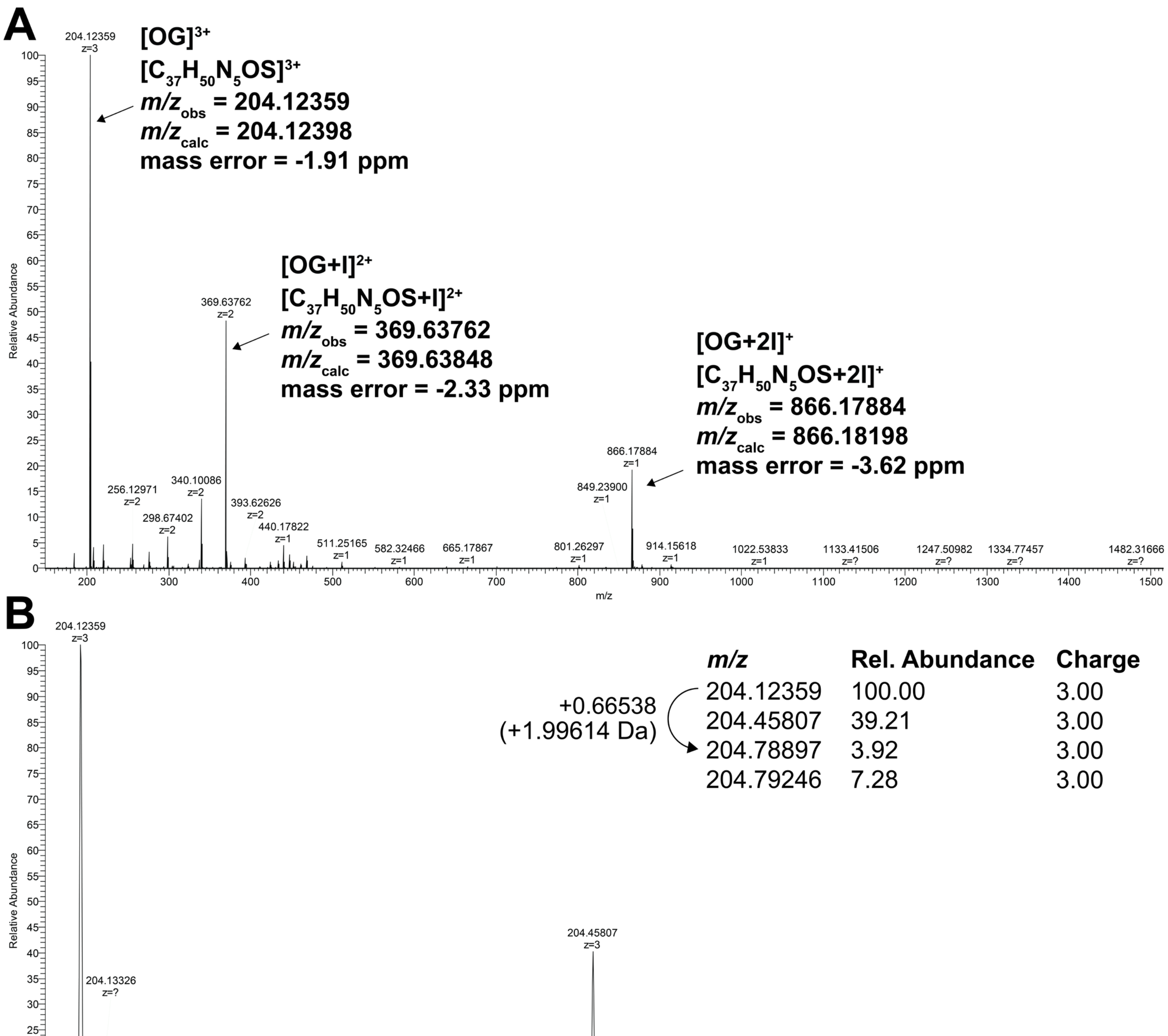


**Figure S21**: MS analysis of OliGreen (OG). (A) $ESI^+$ mass spectrum with m/z range of 150.0-1500.0. The observed and calculated m/z ratios of the main species $[OG]^{3+}$ as well as two minor species ($[OG+I]^{2+}$ and $[OG+2I]^{+}$) are shown. (B) Zoom of the $ESI^+$ mass spectrum (A) showing the isotopic distribution of the main species $[OG]^{3+}$. m/z ratios, relative abundances, and charges of observed peaks are as listed. The characteristic isotopic distribution confirms the presence of exactly one sulfur atom in OG. The mass difference between the monoisotopic peak at m/z 204.12359 and the 2$^{nd}$ isotope peak at m/z 204.78897 is 1.99614 Da, the mass difference between monoisotopic mass of $^{32}S$ (31.97207 Da) and $^{34}S$ (33.96786 Da) is 1.99579 Da. The natural abundance of $^{34}S$ (4.37%) matches the relative abundance of the observed 2$^{nd}$ isotope peak (3.92%).

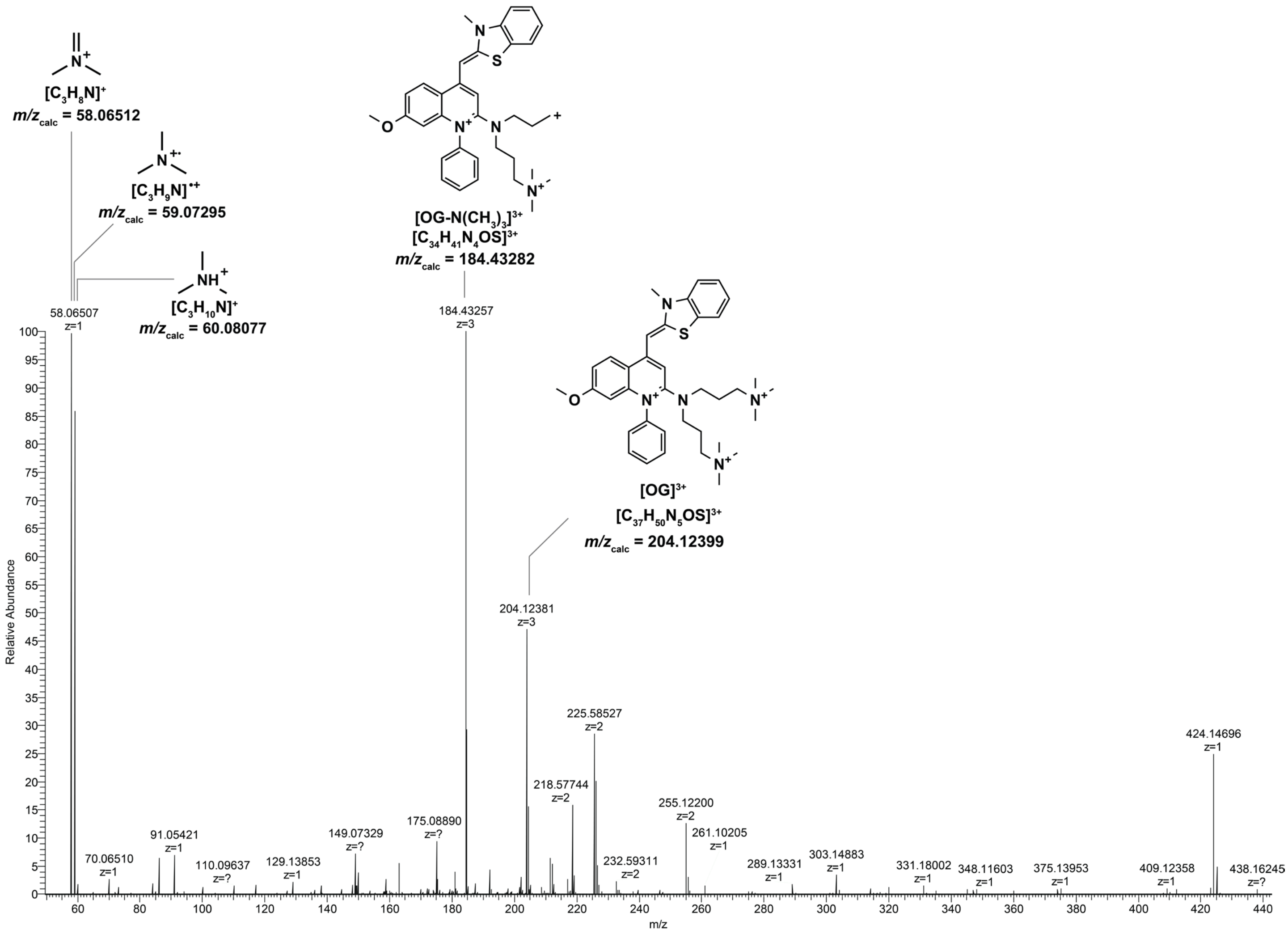


**Figure S22**: MS/MS analysis of OliGreen (OG). The tandem MS spectrum (204.0 at 40% HCD) with m/z range of 50.0-438.2 shows the presence of several fragments of the precursor species $[OG]^{3+}$. Structures, empirical formulae, and calculated m/z ratios are shown for the precursor peak (m/z 204.12381) as well as select fragment peaks.

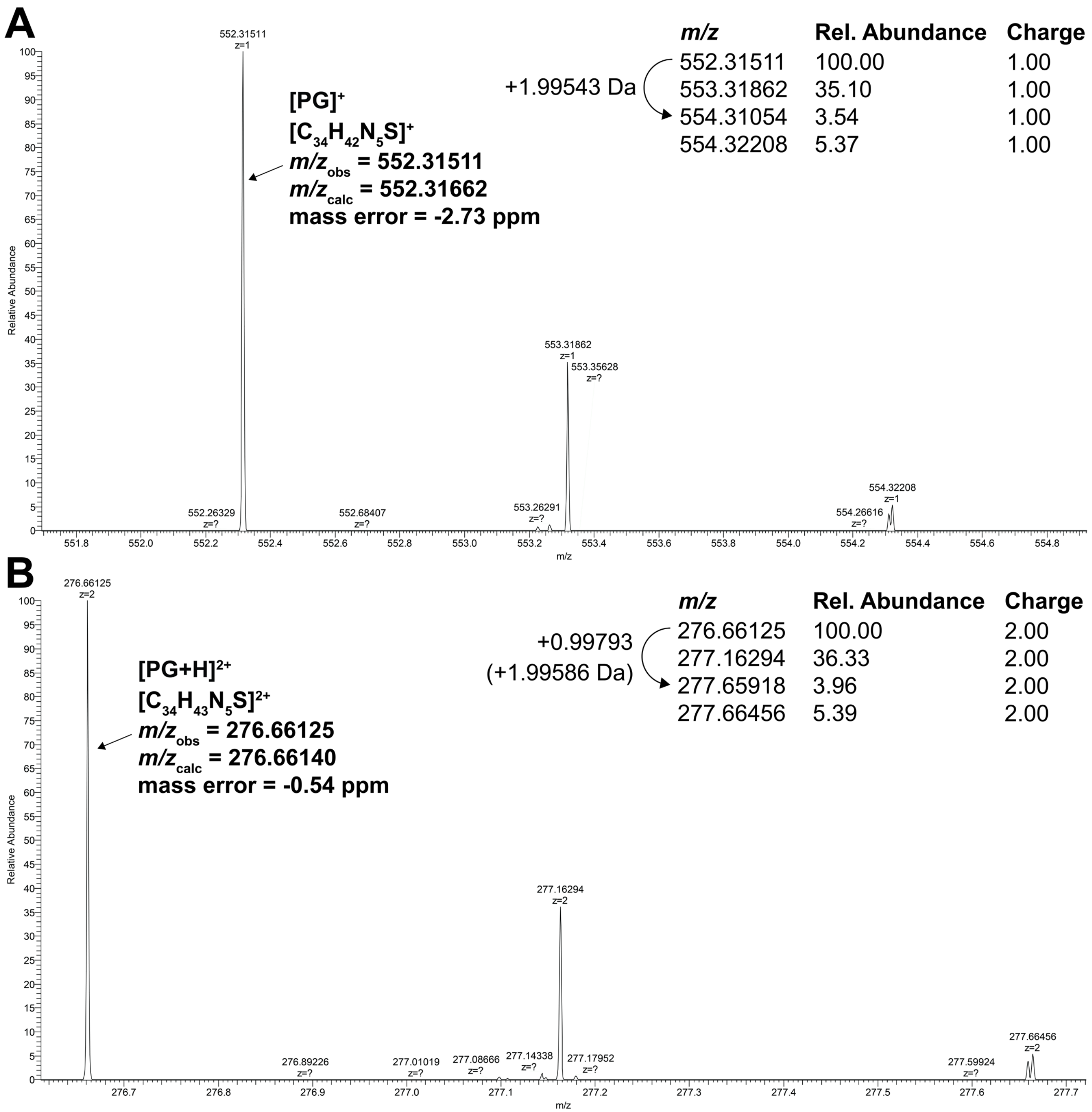


**Figure S23**: MS analysis of PicoGreen (PG). (A) Zoom of the ESI$^+$ mass spectrum (full range of m/z 100.0-1500.0) showing the isotopic distribution of the main species $[PG]^+$ at retention time 13.10-13.55 min. (B) Zoom of the ESI$^+$ mass spectrum (full range of m/z 100.0-1500.0) showing the isotopic distribution of the main species $[PG+H]^{2+}$ at retention time 13.26-13.44 min. The observed and calculated m/z ratios of the main species and m/z ratios and relative abundance of isotopic distribution are as indicated. The characteristic isotopic distributions for both species confirm the presence of exactly one sulfur atom in PG. The mass difference between monoisotopic peak of $[PG]^+$ at m/z 552.31511 and the 2$^{nd}$ isotope peak at m/z 554.31054 is 1.99543, the mass difference between monoisotopic peak of $[PG+H]^{2+}$ at m/z 276.66125 and the 2$^{nd}$ isotope peak at m/z 277.65918 is 1.99586. The mass difference between monoisotopic mass of $^{32}S$ (31.97207 Da) and $^{34}S$ (33.96786 Da) is 1.99579 Da. The natural abundance of $^{34}S$ (4.37%) matches the relative abundance of the observed 2$^{nd}$ isotope peak (3.54% and 3.96% in A and B, respectively).

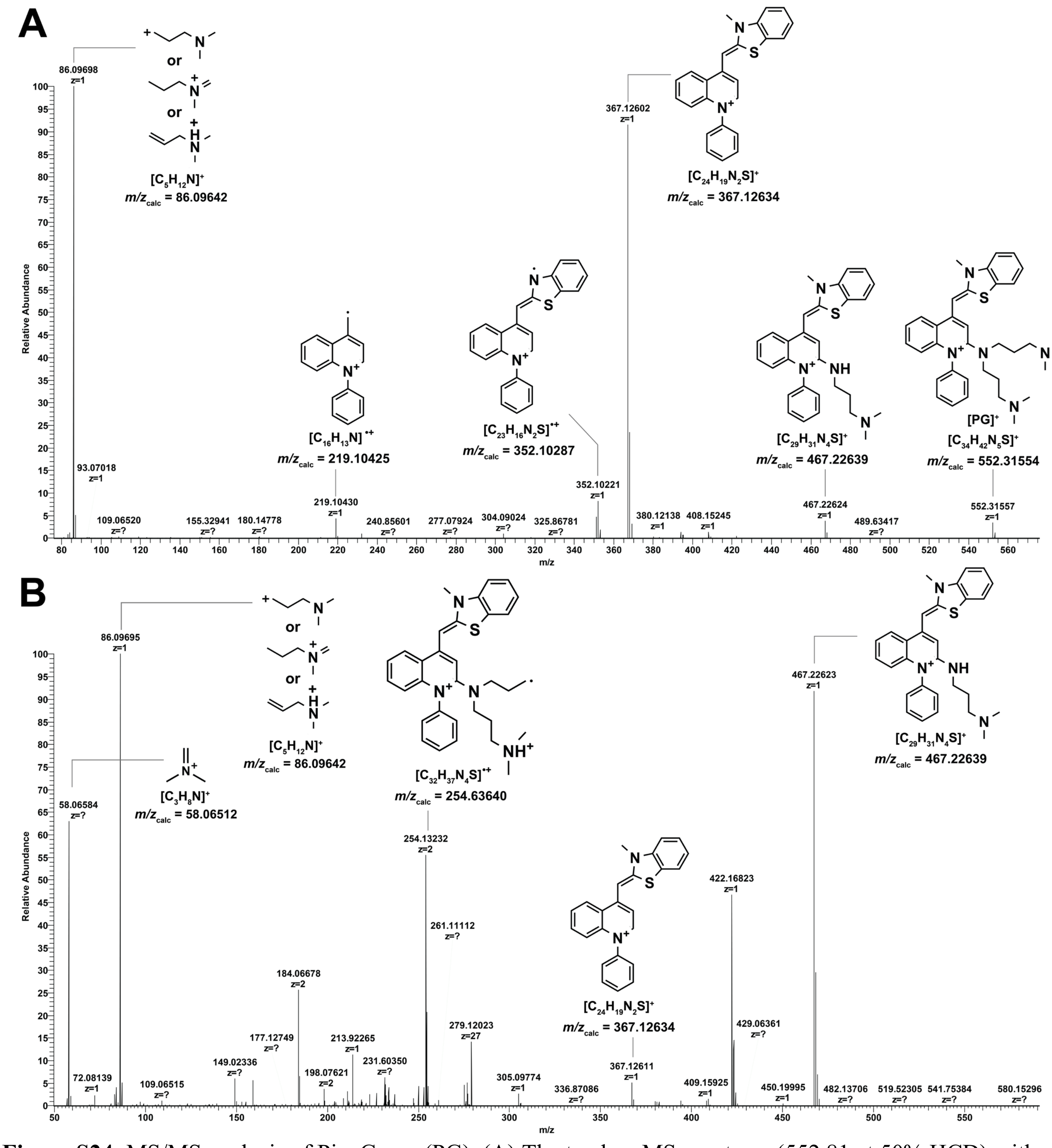


**Figure S24**: MS/MS analysis of PicoGreen (PG). (A) The tandem MS spectrum (552.81 at 50% HCD) with m/z range of 76.67-1150 shows the presence of several fragments of the precursor species $[PG]^+$. Structures, empirical formulae, and calculated m/z ratios are shown for the precursor peak (m/z 552.31557) as well as select fragment peaks. (B) The tandem MS spectrum (552.81 at 30% HCD) with m/z range of 50.0-585.0 shows the presence of several fragments of the precursor species $[PG+H]^{2+}$. Structures, empirical formulae, and calculated m/z ratios are shown for select fragment peaks. The precursor is almost fully depleted and the precursor peak (expected at around m/z 276.66) is weak or absent.

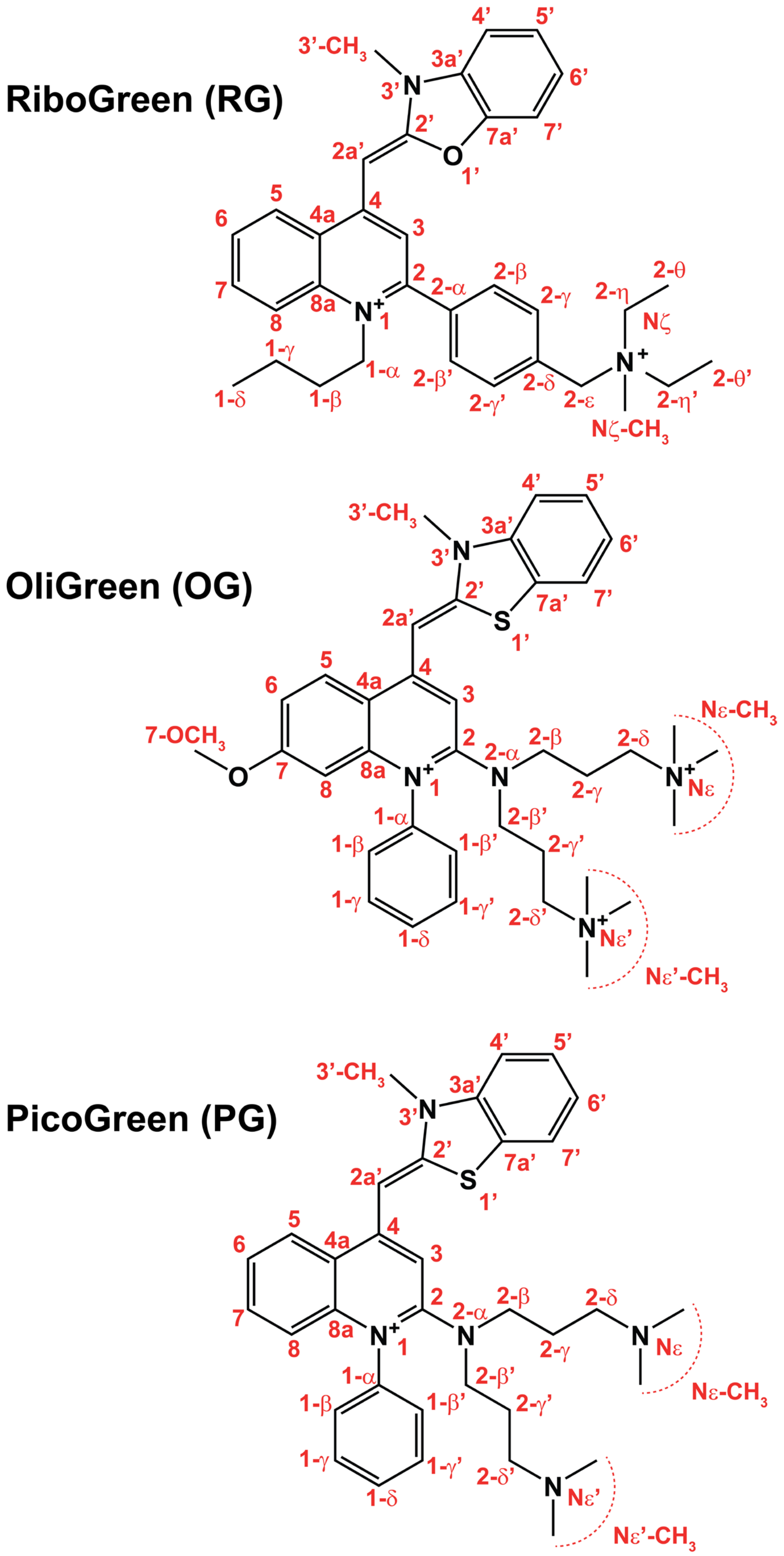


**Figure S25:** The molecular structures of RiboGreen, OliGreen and PicoGreen. Atom numbering is shown.

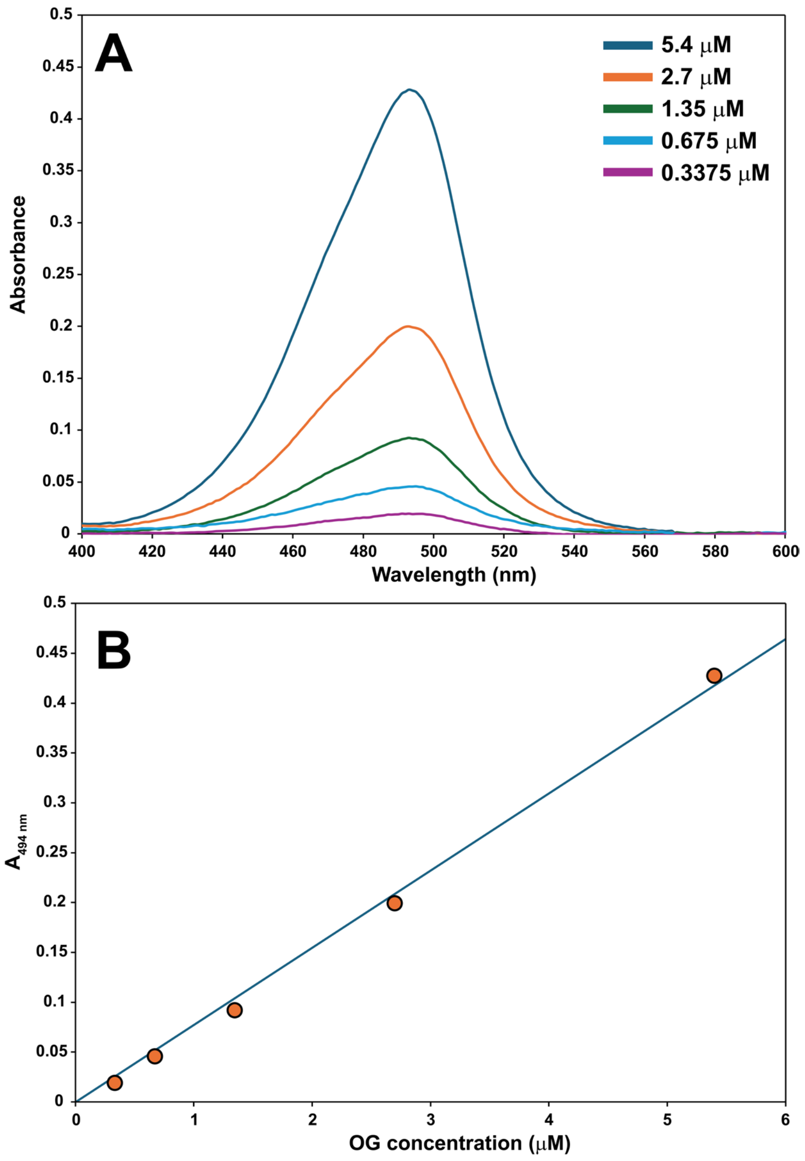


**Figure S26:** Extinction coefficient of OG. (A) Absorption spectra of OG at various concentrations in TE buffer at pH 7.5. The absorption maximum of the free dye in these conditions was found to be at 494 nm. (B) A plot of the absorption at 494 nm ($A_{494\ nm}$) against OG concentration. The extinction coefficient was determined by the slope of the linear fit (blue).

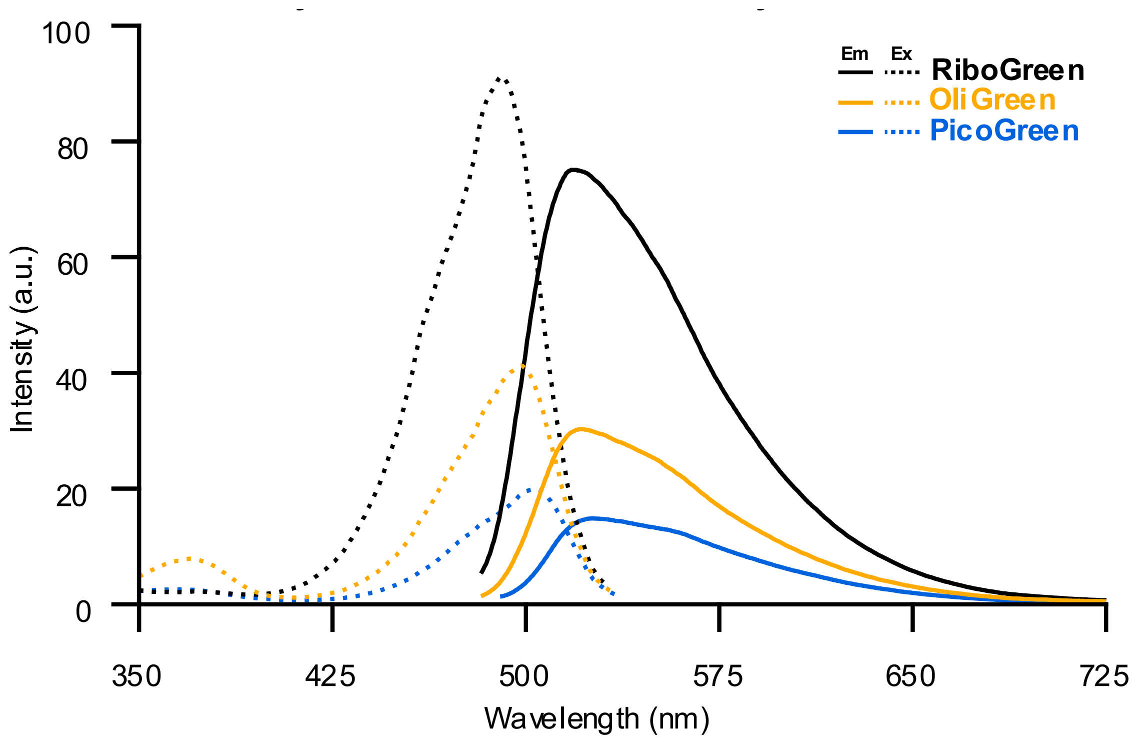


**Figure S27:** Excitation and emission spectra of RG, OG and PG when dissolved in 100% glycerol in the absence of any nucleic acids. The dyes exhibit negligible fluorescence in aqueous environments, absent of nucleic acids. Fluorescence can be recovered by placing the dyes in a high viscosity environment. The final concentration of each dye in the glycerol is a 200-fold dilution from the stock solutions.

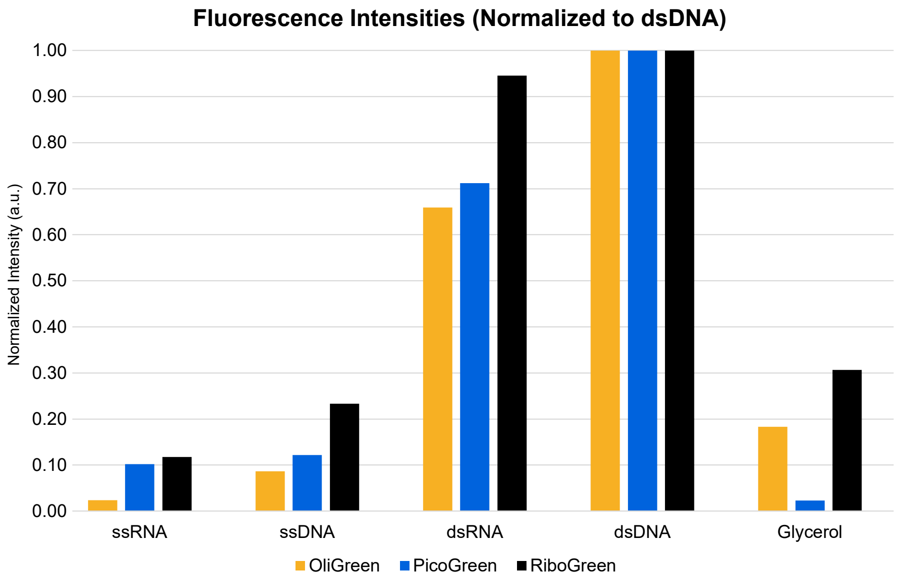


**Figure S28:** A comparison of the maximum fluorescence intensities of RG, OG and PG recorded during their respective emission scans in every condition tested. The intensities are normalized to the intensity of the max signal when bound to dsDNA. All dyes exhibit the strongest fluorescence when bound to dsDNA. Non-normalized intensities are reported in Table S2.

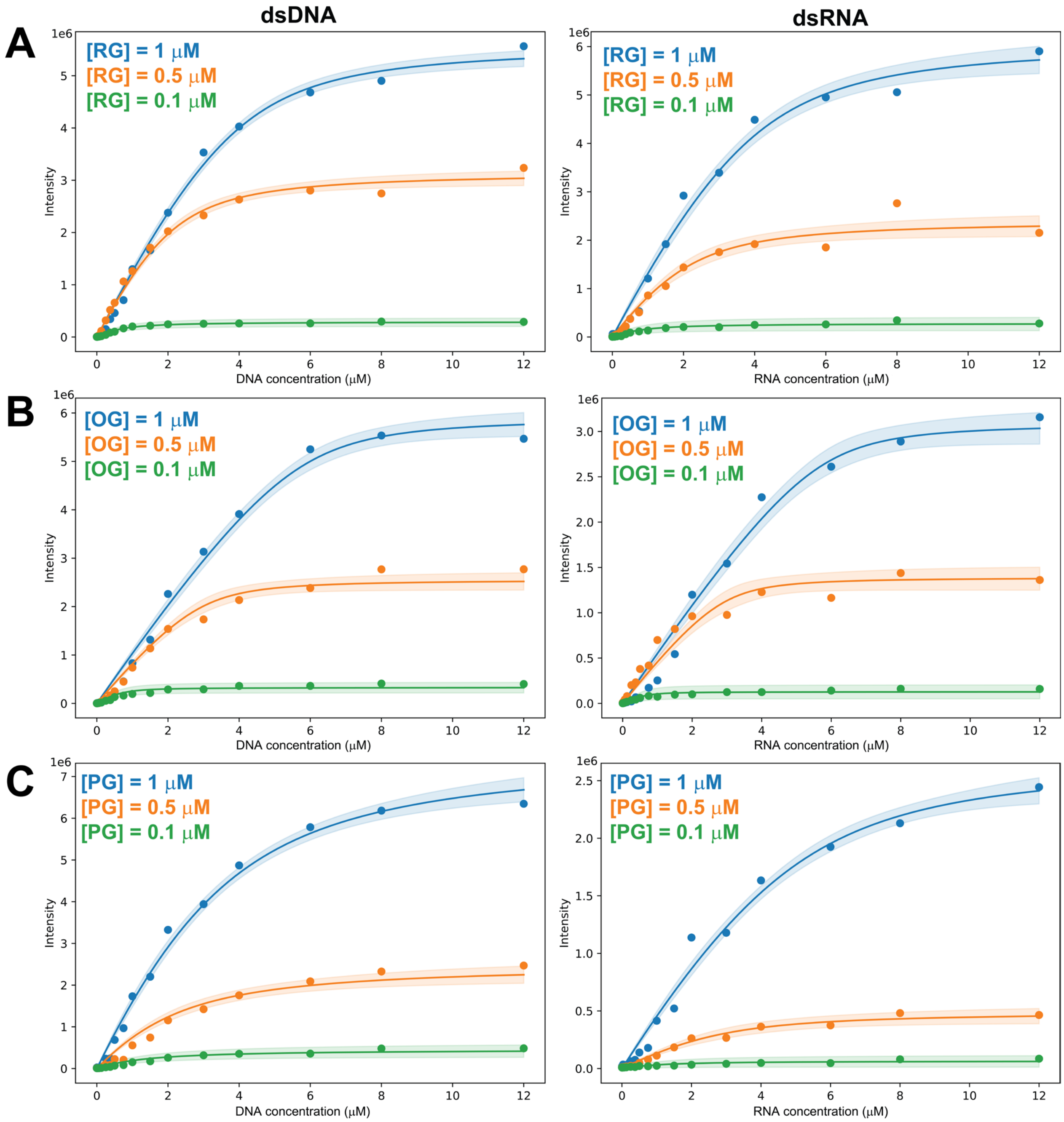


**Figure S29**: Plots of fluorescence intensity versus nucleic acid base pair concentration for (A) RiboGreen, (B) OliGreen, and (C) PicoGreen in the presence of double-stranded DNA (left) and double-stranded RNA (right) at dye concentrations of 1 μM (blue), 0.5 μM (orange), and 0.1 μM (green). Dissociation constants ($K_d$) and binding site size (n) were determined by global fit using the McGhee-von Hippel model (Table 2).

### Supporting Tables

**Table S1:** $^{1}H$ and $^{13}C$ chemical shifts for commercially available unsymmetric cyanine dyes with known chemical shift assignments. The chemical shifts of RG, OG, and PG were determined in this work and the chemical shifts of TO, SSA, SGI, SGII, and SGO were determined previously.[1-3]

| ring system/ | position | Thiazole Orange | | SYBR Safe | | SYBR Green I | | SYBR Green II | | SYBR Gold | | RiboGreen | | OliGreen | | PicoGreen (DMSO) | | PicoGreen (MeOD) | |
|---|---|---|---|---|---|---|---|---|---|---|---|---|---|---|---|---|---|---|---|
| | | $^{1}H$ | $^{13}C$ | $^{1}H$ | $^{13}C$ | $^{1}H$ | $^{13}C$ | $^{1}H$ | $^{13}C$ | $^{1}H$ | $^{13}C$ | $^{1}H$ | $^{13}C$ | $^{1}H$ | $^{13}C$ | $^{1}H$ | $^{13}C$ | $^{1}H$ | $^{13}C$ |
| | | X=S | | X=S | | X=S | | X=O | | X=O | | X=O | | X=S | | X=S | | X=S | |
| benzazolium | 2’ | | 159.7 | | 160.0 | | 158.9 | | 162.1 | | 161.8 | | 162.0 | | 159.7 | | - | | 160.3 |
| | 2a’ | 6.93 | 87.8 | 6.94 | 88.0 | 6.78 | 87.2 | 6.33 | 73.9 | 6.21 | 74.1 | 6.38 | 75.0 | 6.87 | 88.0 | 6.94 | 88.1 | 6.94 | 87.6 |
| | 3’-$CH_3$ | 4.01 | 33.7 | 4.03 | 33.7 | 3.95 | 33.9 | 3.93 | 31.1 | 3.91 | 31.1 | 3.91 | 31.2 | 4.03 | 34.3 | 4.06 | 34.4 | 4.06 | 32.8 |
| | 3a’ | | 140.4 | | 140.4 | | 141.2 | | 131.8 | | 131.9 | | 131.8 | | 141.2 | | - | | 140.8 |
| | 4’ | 7.77 | 112.8 | 7.80 | 112.9 | 7.70 | 112.9 | 7.69 | 111.3 | 7.63 | 111.1 | 7.67 | 111.4 | 7.78 | 113.2 | 7.82 | 113.4 | 7.72 | 112.3 |
| | 5’ | 7.61 | 128.0 | 7.63 | 128.1 | 7.59 | 128.7 | 7.52 | 126.5 | 7.47 | 126.4 | 7.49 | 126.5 | 7.64 | 128.7 | 7.66 | 128.7 | 7.67 | 128.3 |
| | 6’ | 7.41 | 124.3 | 7.43 | 124.4 | 7.39 | 124.6 | 7.44 | 124.9 | 7.33 | 124.5 | 7.35 | 124.9 | 7.44 | 124.8 | 7.46 | 124.9 | 7.46 | 124.5 |
| | 7’ | 8.04 | 122.8 | 8.06 | 122.8 | 7.97 | 123.2 | 7.80 | 111.4 | 7.65 | 111.2 | 7.66 | 111.4 | 8.06 | 123.3 | 8.08 | 123.4 | 8.01 | 122.5 |
| | 7a’ | | 123.7 | | 123.8 | | 123.8 | | 146.5 | | 146.4 | | 146.5 | | 124.0 | | - | | 123.7 |
| 4-quinolinium | 2 | 8.61 | 145.0 | 8.64 | 144.4 | | 158.5 | | 158.1 | | 152.0 | | 153.3 | | 157.8 | | - | | 158.0 |
| | 3 | 7.36 | 107.7 | 7.39 | 107.7 | 7.04 | 102.8 | 8.00 | 106.2 | 7.85 | 112.0 | 7.76 | 112.0 | 7.05 | 102.3 | 7.09 | 103.3 | 7.25 | 102.7 |
| | 4 | | 148.5 | | 148.5 | | 149.1 | | 148.4 | | 148.5 | | 149.4 | | 150.0 | | - | | 150.5 |
| | 4a | | 123.9 | | 124.2 | | 122.0 | | 121.7 | | 124.9 | | 123.8 | | 116.8 | | - | | 122.8 |
| | 5 | 8.80 | 125.4 | 8.81 | 125.7 | 8.61 | 125.5 | 8.76 | 126.1 | 8.02 | 106.8 | 8.84 | 126.6 | 8.71 | 128.0 | 8.75 | 125.7 | 8.61 | 124.5 |
| | 6 | 7.78 | 126.8 | 7.76 | 126.7 | 7.58 | 126.1 | 7.65 | 126.4 | | 158.3 | 7.80 | 127.1 | 7.32 | 114.7 | 7.66 | 126.3 | 7.65 | 125.9 |
| | 7 | 8.02 | 133.1 | 7.99 | 133.2 | 7.67 | 133.0 | 7.74 | 133.6 | 7.75 | 123.6 | 8.05 | 134.1 | | 162.9 | 7.75 | 133.3 | 7.69 | 132.5 |
| | 8 | 8.05 | 118.2 | 8.17 | 118.1 | 7.13 | 119.0 | 6.83 | 118.7 | 8.19 | 121.4 | 8.23 | 119.5 | 6.39 | 102.3 | 7.11 | 119.3 | 7.21 | 118.9 |
| | 8a | | 137.9 | | 137.0 | | 140.7 | | 141.2 | | 134.8 | | 138.5 | | 142.8 | | - | | 140.8 |
| 4-quinolinium substituents ($R^1$, $R^2$, $R^6$, $R^7$) | 1-α | 4.17 | 42.3 | 4.57 | 55.4 | | | | | 3.94 | 40.4 | 4.35 | 49.9 | | 138.9 | | - | | 138.8 |
| | 1-β/β‘ | | | 1.89 | 22.1 | - | - | - | - | - | - | 1.70 | 30.9 | 7.75 | 129.8 | 7.74 | 130.1 | 7.64 | 129.5 |
| | 1-γ/γ‘ | | | 0.96 | 10.6 | - | - | - | - | - | - | 1.14 | 19.4 | 7.84 | 131.2 | 7.81 | 131.1 | 7.83 | 130.6 |
| | 1-δ | | | | | | | | | | | 0.65 | 13.5 | 7.74 | 131.0 | 7.73 | 131.0 | 7.77 | 130.2 |
| | 2-α | | | | | | | | | | 137.1 | | 137.4 | | | | | | |
| | 2-β/β‘ | | | | | 3.26/3.16 | 50.4/54.8 | 3.49 | 31.8 | 7.88 | 130.1 | 7.88 | 129.7 | 3.21 | 49.5 | 3.24 | 49.6 | 3.37 | 49.8 |
| | 2-γ/γ‘ | | | | | 1.49/1.23 | 24.7/20.3 | 2.71 | 56.8 | 7.84 | 134.1 | 7.86 | 134.1 | 1.72 | 20.9 | 1.70 | 22.0 | 1.76 | 23.2 |
| | 2-δ/δ‘ | | | | | 2.19/0.73 | 56.3/11.5 | | | | 130.5 | | 130.5 | 3.15 | 63.3 | 2.89 | 54.8 | 2.74 | 55.5 |
| | 2-ε | | | | | | | | | 4.64 | 63.5 | 4.65 | 63.5 | | | | | | |
| | Nε/ε‘-$CH_3$ | | | | | 2.12 | 45.2 | | | | | | | 3.03 | 53.0 | 2.75 | 42.9 | 2.61 | 43.1 |
| | Nδ-$CH_3$ | | | | | | | 2.20 | 45.3 | | | | | | | | | | |
| | Nζ-$CH_3$ | | | | | | | | | 2.97 | 46.7 | 2.96 | 46.8 | | | | | | |
| | 2-η/η‘ | | | | | | | | | 3.33/3.43 | 55.8 | 3.32/3.41 | 56.0 | | | | | | |
| | 2-θ/θ‘ | | | | | | | | | 1.38 | 8.9 | 1.38 | 8.2 | | | | | | |
| | 6-$OCH_3$ | | | | | | | | | 4.08 | 56.8 | | | | | | | | |
| | 7-$OCH_3$ | | | | | | | | | | | | | 3.73 | 56.2 | | | | |

**Table S2:** The fluorescence intensity maximum and wavelength recorded on each dye in every condition tested. Intensity values are the raw output from the instrument. Wavelengths are rounded to the nearest nm.

| Dye | Condition | $A_{max}$ (nm) | Emission Intensity at $A_{max}$ |
|---|---|---|---|
| OliGreen | ssDNA | 523 | 14.27 |
| OliGreen | dsDNA | 518 | 165.45 |
| OliGreen | ssRNA | 523 | 3.97 |
| OliGreen | dsRNA | 523 | 109.00 |
| OliGreen | 100% Glycerol | 522 | 30.35 |
| PicoGreen | ssDNA | 526 | 78.88 |
| PicoGreen | dsDNA | 523 | 648.79 |
| PicoGreen | ssRNA | 527 | 66.11 |
| PicoGreen | dsRNA | 526 | 462.13 |
| PicoGreen | 100% Glycerol | 527 | 14.88 |
| RiboGreen | ssDNA | 528 | 57.06 |
| RiboGreen | dsDNA | 523 | 244.73 |
| RiboGreen | ssRNA | 523 | 28.80 |
| RiboGreen | dsRNA | 524 | 231.31 |
| RiboGreen | 100% Glycerol | 520 | 74.94 |

### Supporting Files

**DNA_RG_1.tar.gz: MD simulation of RG and double-stranded DNA (Run 1)**

DNA_RG_1.tar.gz is a gzipped archive containing the starting structure DNA_RG_1.pdb and the trajectory file DNA_RG_1.dcd. This run is comprised of 21,750 frames.

**DNA_RG_2.tar.gz: MD simulation of RG and double-stranded DNA (Run 2)**

DNA_RG_2.tar.gz is a gzipped archive containing the starting structure DNA_RG_2.pdb and the trajectory file DNA_RG_2.dcd. This run is comprised of 9,858 frames.
